# Sex-Specific Effects of Anxiety on Cognition and Activity-Dependent Neural Networks: Insights from (Female) Mice and (Wo)Men

**DOI:** 10.1101/2023.07.07.548180

**Authors:** Holly C. Hunsberger, Seonjoo Lee, Michelle Jin, Marcos Lanio, Alicia Whye, Jiook Cha, Miranda Scarlata, Keerthana Jayaseelan, Christine. A. Denny

## Abstract

**INTRODUCTION:** Neuropsychiatric symptoms (NPS), such as depression and anxiety, are observed in 90% of Alzheimer’s disease (AD) patients, two-thirds of whom are women. NPS usually manifest long before AD onset creating a therapeutic opportunity. Here, we examined the impact of anxiety on AD progression and the underlying brain-wide neuronal mechanisms.

**METHODS:** To gain mechanistic insight into how anxiety impacts AD progression, we performed a cross-sectional analysis on mood, cognition, and neural activity utilizing the ArcCreER^T2^ x enhanced yellow fluorescent protein (eYFP) x APP/PS1 (AD) mice. The ADNI dataset was used to determine the impact of anxiety on AD progression in human subjects.

**RESULTS:** Female AD mice exhibited anxiety-like behavior and cognitive decline at an earlier age than control (Ctrl) mice and male mice. Brain-wide analysis of c-Fos^+^ revealed changes in regional correlations and overall network connectivity in AD mice. Sex-specific memory trace changes were observed; female AD mice exhibited impaired memory traces in dorsal CA3 (dCA3), while male AD mice exhibited impaired memory traces in the dorsal dentate gyrus (dDG). In the ADNI dataset, anxiety predicted transition to dementia. Female subjects positive for anxiety and amyloid transitioned more quickly to dementia than male subjects.

**CONCLUSIONS:** While future studies are needed to understand whether anxiety is a predictor, a neuropsychiatric biomarker, or a comorbid symptom that occurs during disease onset, these results suggest that AD network dysfunction is sexually dimorphic, and that personalized medicine may benefit male and female AD patients rather than a one size fits all approach.

## INTRODUCTION

Alzheimer’s disease (AD), a progressive and debilitating neurodegenerative and psychiatric disorder, stands alone as one of the ten leading causes of death in the United States that cannot be prevented, slowed, or cured (1). Neuropsychiatric symptoms (NPS), such as depression and anxiety, are observed in 90% of AD patients and symptoms usually manifest long before AD onset (2). Although much of the field has focused on the link between depression and AD, recent clinical evidence supports that anxiety can predict the progression to AD more than depression, brain atrophy, and cognitive impairment (3). *The Harvard Aging Brain Study* recently reported that higher amyloid burden was associated with anxiety over time in cognitively normal older individuals (4), meaning NPS represent a potential biomarker of AD.

Females represent two-thirds of the AD population and are more susceptible to depression and anxiety; however, the majority of preclinical AD studies have focused primarily on male subjects (1). It was once assumed that women’s susceptibility was due to longer lifespans, but statistics now demonstrate that the lifespan for men and women is similar, differing by only 4-5 years (4). Interestingly, in the APP695SWE AD mouse model, plaque burden is three times greater in female mice than males, indicating disease progression is not similar in both sexes (5). Therefore, understanding why females are more susceptible to NPS in relation to AD has therapeutic potential.

Recent genetic technologies have allowed for unprecedented access into the mechanisms of memory loss and allowed for single-cell resolution of individual memory traces or engrams. We and others have utilized activity-dependent tagging murine lines such as the ArcCreER^T2^ x eYFP mice (6), to better understand, where, when, and how AD impacts individual memories. We have previously shown that deficits in contextual fear memory in male AD mice are associated with a decrease in memory trace reactivation in the dentate gyrus (DG), and that optogenetic stimulation of the neurons active during encoding can restore memory retrieval (9). However, recent activity-dependent studies have not extensively included females, nor investigated how anxiety influences these memory traces.

Considering the clinical evidence that female AD patients exhibit greater anxiety compared to males, we sought to determine whether the onset of anxiety-like behavior was correlated with cognitive decline and the corresponding neural ensembles in a sex-specific manner. Ctrl and AD mice were administered behavioral assays to quantify mood and behavior. We report that female AD mice exhibited anxiety-like behavior at an earlier age when compared to Ctrl mice. Female AD mice additionally displayed memory deficits as early as 2 months of age, whereas male AD mice displayed memory deficits at 6 months of age. The increased anxiety-like behavior correlated with cognitive decline only in AD female mice. Female AD mice showed altered memory traces in dCA3 and an uncorrelated brain-wide neural network, unlike males. To validate these preclinical findings, we analyzed the impact of anxiety on AD in human subjects using the Alzheimer’s disease neuroimaging initiative (ADNI) dataset. Female AD subjects: 1) exhibited higher anxiety; 2) with amyloid deposition transitioned faster to dementia; and 3) with anxiety had smaller brain volumes. Lastly, we found that anxiety is the most significant predictor of dementia transition. These data suggest that anxiety impacts cognitive decline and AD progression is sex-specific with brain-wide network changes specifically in females.

## MATERIALS AND METHODS

Full description of Methods and Materials available in Supplement 1.

### Mice

The amyloid precursor protein/presenilin 1 (APP/PS1) mice (12) were previously bred with the ArcCreER^T2^ (8) x channelrhodopsin (ChR2)-enhanced yellow fluorescent protein (eYFP) mice (9) as previously described (7). Female and male mice were used in all experiments. Mice were group housed (4-5/cage) in a 12-h light/dark cycle (lights on at 0600 hours) colony room maintained at 22°C. Mice had *ad libitum* access to food and water. All procedures were conducted in accordance with the National Institutes of Health regulations and by the Institutional Animal Care and Use Committee (IACUC) of the New York State Psychiatric Institute (NYSPI).

### Alzheimer’s Disease Neuroimaging Initiative (ADNI)

Data used in the preparation of this article were obtained from the ADNI database (adni.loni.usc.edu). ADNI was launched in 2003 as a public-private partnership, led by Michael W. Weiner, M.D (10). The primary goal of ADNI has been to test whether serial magnetic resonance imaging (MRI), positron emission tomography (PET), other biological markers, and clinical and neuropsychological assessment can be combined to measure the progression of mild cognitive impairment (MCI) and early AD. The dataset used was downloaded on October 28^th^, 2018.

## RESULTS

### Female AD mice exhibit anxiety-like behavior and cognitive decline at an earlier age than male AD mice

Although there are numerous studies on cognitive deficits, there are few studies on mood abnormalities in APP/PS1 mice (11–18). There is conflicting literature on behavioral performance, and it is often difficult to compare between studies because of background strain differences. Therefore, we first characterized mood and cognition at 2, 4, and 6 months of age in both female and male Ctrl and APP/PS1 (AD) mice on a 129S6/SvEv background strain (7) (**Fig. 1A, Table S1**). In the open field (OF), all groups had a comparable center distance (%) (**Fig. 1B**). Locomotion was similar in all groups (**Fig. S1A**). In the elevated plus maze (EPM), female mice spent less time in the open arms with age, starting at 4 months of age (**Fig. 1C, Fig. S1B**). In the marble burying (MB) assay, a measure of perseverative behavior (19), 4-month-old female AD mice buried more marbles (**Fig. 1D**). In the light dark test (LDT), female AD mice exhibited decreased activity on the light side, as early as 2 months of age when compared to Ctrl mice who also displayed an age-dependent increase in anxiety (**Fig. 1E, S1C-S1D**). In the novelty suppressed feeding (NSF) paradigm, a measure of hyponeophagia, both female Ctrl and AD mice exhibited comparable behaviors (**Fig. 1F-1G**).

**Fig. 1.**
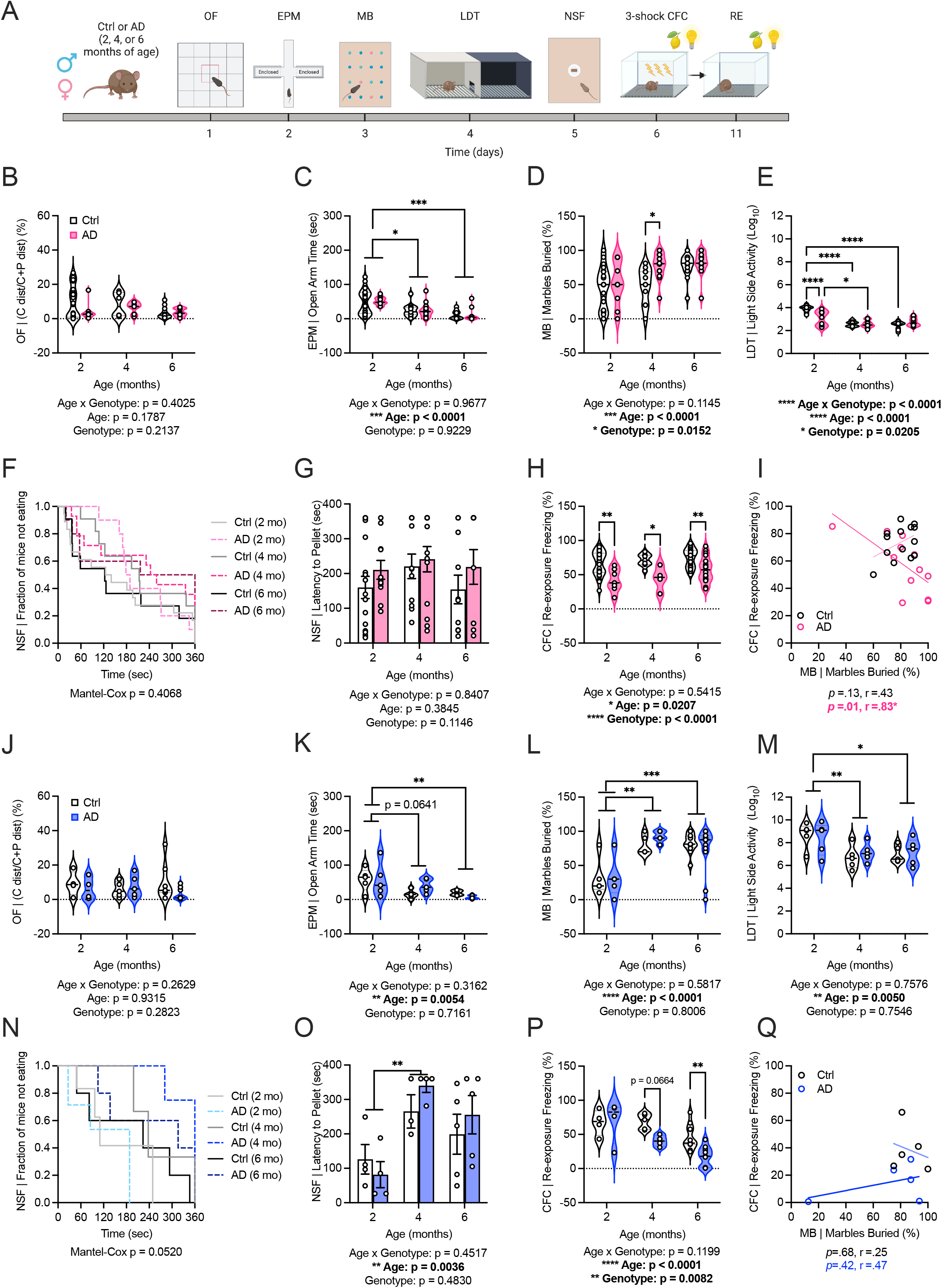
Female AD mice exhibit earlier anxiety-like behavior and cognitive decline. (**A**) Experimental timeline. (**B**) Ctrl and AD female mice exhibit a similar percent distance in the center of the OF. (**C**) Female Ctrl and AD mice spend less time in the open arms of the EPM with age. (**D**) Female AD mice bury a larger percentage of marbles at 4 months of age compared to Ctrl mice. (**E**) Female AD mice spend less activity on the light side of the box at 2 months of age compared to Ctrl mice. Female Ctrl and AD mice display less activity on the light side with age. (**F-G**) In the NSF, latency to the pellet is similar between female Ctrl and AD mice. **(H)** Female AD mice exhibit cognitive deficits at 2 months of age compared to Ctrl mice in the CFC test. (**I**) Freezing (%) is correlated with increased marbles buried in 6-month-old female AD mice. (**J**) Male Ctrl and AD mice exhibit similar percent distance in the center of the OF. (**K**) Male AD mice spend less time in the open arms of the EPM with age. (**L**) Male Ctrl and AD male mice bury more marbles with age. **(M)** Age significantly impacts time spent on the light side in the LDT. **(N-O)** Latency time to the food pellet is similar in all groups when graphed in a survival plot; however; Age influence latency to eat in the RMANOVA. (**P**) Male AD mice are impaired in CFC freezing (%) at 6 months of age. (**Q**) Freezing (%) is not correlated with increased marbles buried in 6-month-old male AD mice. (n=4-18 mice per group). Error bars represent ± SEM. *p<0.05, **p<0.01, ***p<0.001. OF, open field; EPM, elevated plus maze; MB, marble burying; LDT, light dark test; NSF, novelty suppressed feeding; CFC, contextual fear conditioning; RE, re-exposure; C, center; P, periphery; dist, distance; sec, seconds; Ctrl, control; AD, Alzheimer’s disease.

Cognitive decline was measured using a 3-shock contextual fear conditioning (CFC) paradigm. Female AD mice showed a decrease in freezing (%) as early as 2 months of age (**Fig. 1H, S1E-S1F**). To determine if cognitive performance was related to anxiety-like behavior, correlations were performed. Marbles buried (%) was negatively correlated with freezing (%) in female AD mice at 6 months of age, suggesting a relationship between perseverative and cognitive behaviors (**Fig. 1I**).

In male AD mice, we confirmed our previous finding that AD does not alter behavior in the OF (7) (**Fig. 1J; Fig. S1G**). In the EPM, MB, and LDT paradigms, there was a significant effect of Age, not of Genotype (**Fig. 1K-1M, S1G-S1J**). In the NSF paradigm, latency to eat was comparable across all genotypes, but when two-way ANOVA was performed, there was a significant impact of Age (**Fig. 1N-1O**). Male AD mice did not show cognitive decline until 6 months of age (**Fig. 1P, Fig. S1K-S1L)** similar to our previous study in male AD mice (7). Unlike female mice, there was no significant correlation between marbles buried (%) and freezing (%) at 6 months of age (**Fig. 1Q**). These data suggest that female and male APP/PS1 mice have distinct trajectories of behavioral impairments.

### Increased negative brain-wide correlations and reduced cluster connectivity in female AD mice during memory retrieval

To understand the neural mechanisms underlying these behavioral changes, we utilized a brain-wide clearing and quantification pipeline (**Fig. 2A**) to quantify neural activity using the immediate early gene (IEG) c-Fos^+^ (20,21). Following 3-shock CFC context re-exposure, 6-month-old Ctrl and AD brains were processed and stained for c-Fos^+^ (**Fig. 2B-2C**). Heatmap correlations between brain regions revealed an increase in negative correlations in female AD mice (**Fig. 2D-2E**). Conversely, male AD mice display increased positive correlations (**Fig. 2F-2G**). In female mice, volcano and parallel plots confirmed the pattern observed in the heatmaps; female AD mice exhibit a greater number of decreased interregional correlations relative to Ctrl mice (**Fig. 2H-2I**). Interestingly, the subiculum (Sub) to cortical amygdala (COA) was the only significantly functional connection negatively decreased in female Ctrl mice. Hippocampal amygdala connections are essential for fear memory and this balance may be important for memory retrieval (22,23). In male AD mice, volcano and parallel plots reveal greater number of positive correlations specifically in the hippocampus (HPC) and retrosplenial cortex (RSP) (**Fig. 2J-K**). These regions are heavily involved in fear memory and might be overactive in AD male mice leading to hyperexcitability often observed early in AD pathology (24–26).

**Fig. 2.**
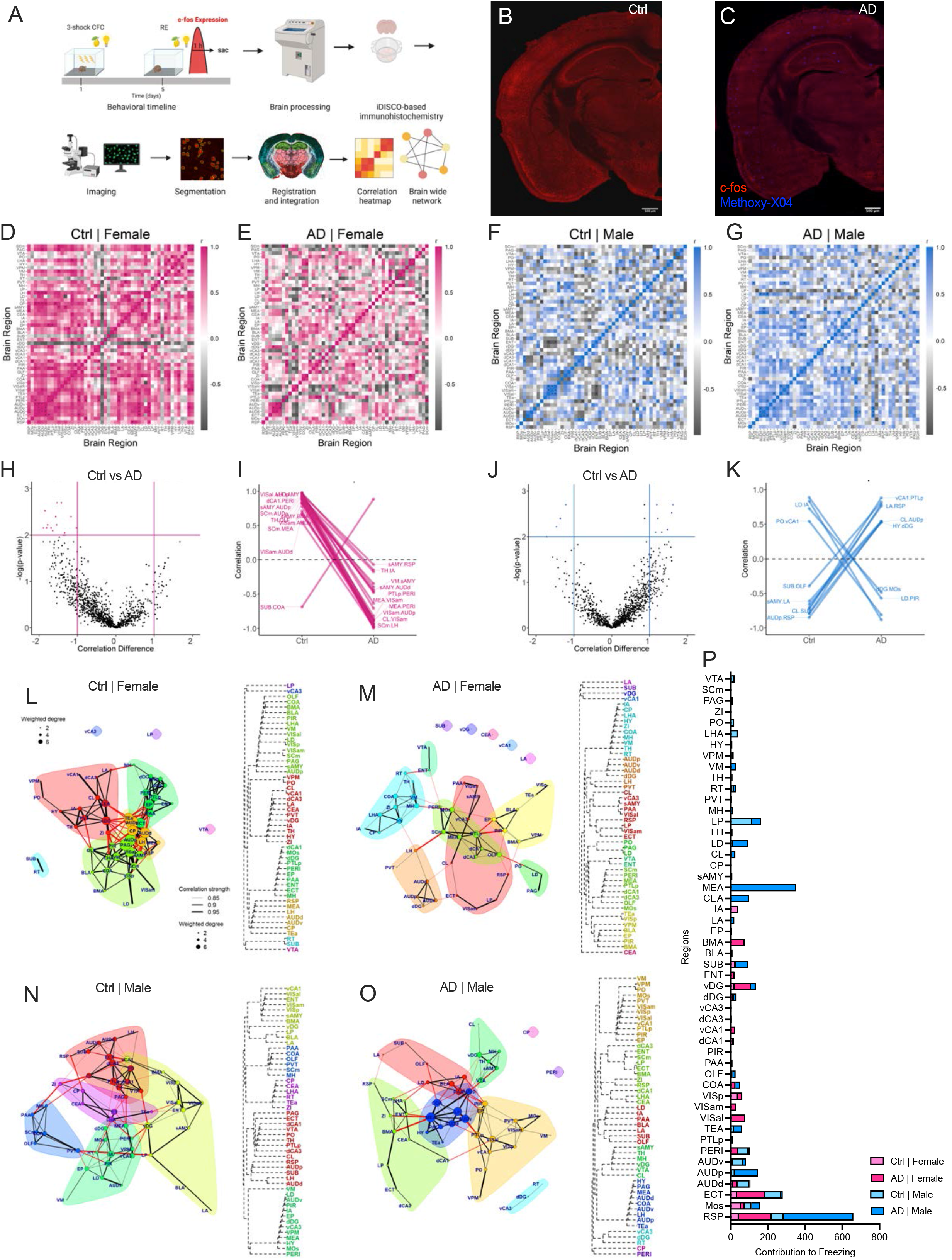
AD alters the correlated activity between brain regions during fear memory retrieval. (**A**) Experimental timeline. (**B-C**) Representative c-Fos^+^ images in Ctrl and AD mice, with Methoxy-X04 staining amyloid plaques. (**D-E**) Brain-wide heat map correlations include the hippocampus, amygdala, thalamus, and hypothalamus. In female mice, pink indicates strong positive correlations while grey indicates strong negative correlations. (**F-G**) Brain-wide heat map correlations include the hippocampus, amygdala, thalamus, and hypothalamus. In male mice, blue indicates strong positive correlations while grey indicates strong negative correlations. (**H**) Volcano plot highlighting a subset of regional correlations with a correlation difference higher than 1 between Ctrl and AD mice. (**I**) Parallel plots confirm the pattern observed in the heat, female AD mice exhibit a greater number of negative correlations compared to positive correlations whereas female Ctrl mice show more positive correlations. (**J-K**) Volcano plot highlighting a subset of regional correlations with a correlation difference higher than 1 between Ctrl and AD mice. Male AD mice display increased positive correlations while male Ctrl mice exhibit increased negative correlations throughout the brain. (**L-M**) Cluster map analysis revealed sparse connectivity in female AD mice, while female Ctrl mice exhibit an integrated network of regions. (**N-O**) The cluster map is highly interconnected in male Ctrl mice, while the cluster map is less cohesive in male AD mice. (**P**) The regions that most contribute to freezing behavior include the RSP, ECT, MEA, vDG, and LP. (n=4-7 mice per group). Error bars represent ± SEM. *p<.05, **p<.01, ***p<.001. CFC, contextual fear conditioning; RE, re-exposure; iDISCO, immunolabeling-enabled imaging of solvent-cleared orgrans; Ctrl, control; AD, Alzheimer’s disease; all other brain regions are defined in supplemental tables.

Cluster network maps were generated to visualize brain regions that clustered as functional communities during CFC retrieval. Analysis revealed the disconnect of the Sub, central amygdala (CEA), ventral CA1 (vCA1), lateral amygdala (LA), and ventral dentate gyrus (vDG) in female AD mice (**Fig. 2L-M**). In male AD mice, there was a disconnect between the caudate putamen (CP), perirhinal cortex (PERI), ventral CA3 (vCA3), dorsal DG (dDG), and the reticular nucleus of the thalamus (RT) (**Fig. 2N-O**). Using a bootstrap prediction analysis, the regions contributing most to freezing behavior were next identified (**Fig. 2P**). The RSP and amygdala (AMG) contributed most in female Ctrl mice whereas the RSP and ectorhinal area (ECT) contributed most in female AD mice. In male mice, the lateral posterior nucleus of the thalamus (LP) and ECT were important for memory retrieval, while the RSP and medial amygdala (MEA) were more prominent in male AD mice, indicating the role of the RSP in memory retrieval.

### Sex-specific changes in brain-wide functional connectivity in AD mice

We next assessed circuit-level connectivity of CFC correlations, modeling the regions as nodes and the correlations as edges (**Fig. 3A-3D, Table S2**). The fear retrieval network was sparser in female AD mice (**Fig. 3A-3B**). In female Ctrl mice, there were strong positive correlations between the HPC and AMG regions. In female AD mice, connections were strongest between dCA1 and vCA1. Interestingly, in both Ctrl and AD mice, the vDG was negatively correlated, but the correlated regions differed. Conversely, male AD mice exhibited networks that were more positively correlated (**Fig. 3C-3D**). However, these positive correlations and connections were lacking in the HPC and instead were overactive in the AMG and auditory regions. The entorhinal cortex (Ent), one of the first regions affected in AD, was negatively correlated in Ctrl mice but positively correlated in AD mice.

**Fig 3.**
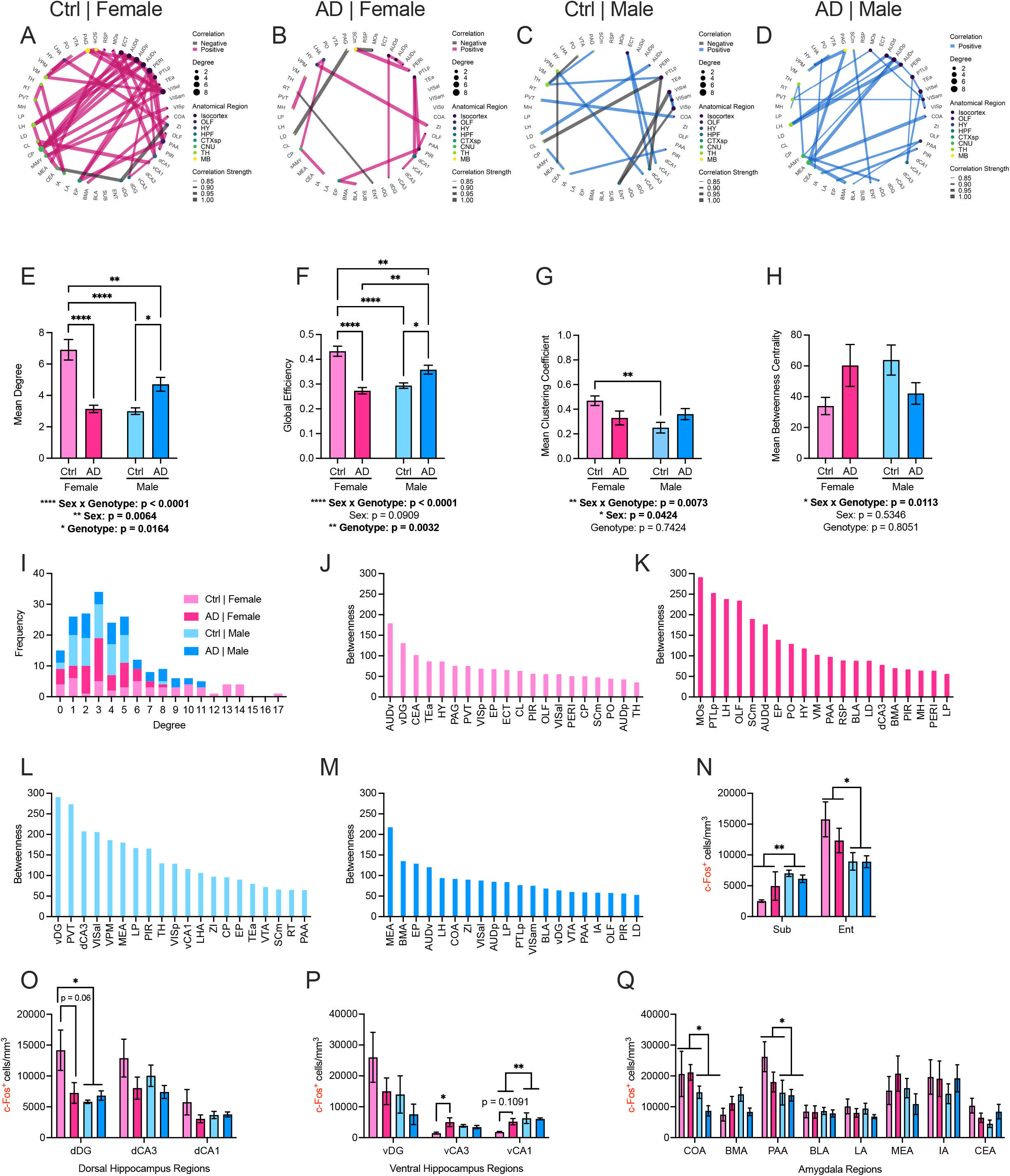
Sex-specific changes in brain-wide functional connectivity during memory retrieval in AD mice. (**A-B**) The fear retrieval network in female AD mice is sparser when compared to female Ctrl mice (grey = negative correlations; pink=positive correlations). Thicker lines indicate stronger correlations and larger circles represent an increased degree of nodes. (**C-D**) In male mice, an opposite pattern is observed. Male AD mice networks are more positively correlated and connected compared to male Ctrl mice. (**E-F**) Male Ctrl and AD mice, and female AD mice exhibit a decrease in mean degree and global efficiency compared to female Ctrl mice. (**G**) Male Ctrl mice have a decreased mean clustering coefficient when compared to female Ctrl mice. (**H**) Mean betweenness centrality is similar among the groups. (**I**) Frequency for each level of degree among the groups. (**J-M**) Betweenness scores for each region reveal the importance of the vDG in females Ctrl and male Ctrl mice. (**N**) Male mice have increased activation in the subiculum whereas female mice show increased c-Fos^+^ activation in the entorhinal cortex. (**O**) In the dHPC, Female Ctrl and AD mice, and male AD mice exhibit decreased c-Fos^+^ activation in the dDG compared to females Ctrl mice. Similar levels are observed in dCA3 and dCA1 between the groups. (**P**) In the vHPC, c-Fos^+^ levels are similar in the vDG but are elevated in female AD mice in vCA3. There is a significant effect of Sex in vCA1. (**Q**) c-Fos^+^ activation is similar throughout the amygdalar regions except in the COA where female mice exhibit increased levels compared to male mice. (n=4-7 mice per group). Error bars represent ± SEM. *p<0.05, **p<0.01, ***p<0.001. Ctrl, control; AD, Alzheimer’s disease; mm, millimeters; Sub, subiculum; Ent, entorhinal cortex; dDG, dorsal dentate gyrus; dCA3, dorsal cornu ammonis 3; dCA1, dorsal cornu ammonis 1; vDG, ventral dentate gyrus; vCA3, ventral cornu ammonis 3; vCA1, ventral cornu ammonis 1; COA, cortical amygdala area; BMA, basomedial amygdala; PAA, piriform-amygdala area; BLA, basolateral amygdala; LA, lateral amygdala; MEA, medial amygdala; IA, intercalated amygdala; CEA, central amygdala; all other brain regions are defined in supplemental tables.

We computed network measures to better understand the structure and properties of these networks (**Fig. 3E-3H**). Female AD mice exhibited a decreased mean degree, whereas male AD mice displayed a greater mean degree compared to Ctrl mice (**Fig. 3E**). A similar pattern emerged with less global efficiency in female AD mice and greater global efficiency in male AD mice (**Fig. 3F**). For clustering coefficient measures, female Ctrl mice exhibited greater clustering when compared to male Ctrl mice (**Fig. 3G**). There was a Sex x Genotype interaction for mean betweenness centrality, but no post-hoc tests reached significance (**Fig. 3H**). The frequency for each level of degree of connectivity, per condition, revealed the same pattern that female AD mice had a higher frequency at lower degrees while male AD mice showed a greater frequency at higher degrees (**Fig. 3I**). Betweenness values for individual regions indicated the importance of the vDG. In both female and male Ctrl mice, the vDG had a higher betweenness score, while it was absent in both female and male AD mice (**Fig. 3J-M**). This finding led us to examine c-Fos^+^ expression in the HPC.

In the Sub, c-Fos^+^ expression was higher in male mice than in female mice, whereas in the Ent, c-Fos^+^ expression was higher in female mice than in male mice (**Fig. 3N**). In the dHPC, male mice exhibited decreased c-Fos^+^ expression in the dDG (**Fig. 3O**). There was no effect of Sex or Genotype in dCA3 or dCA1. In the vDG, there was no effect of Sex or Genotype. In vCA3, female AD mice exhibited greater c-Fos^+^ expression. In vCA1, c-Fos^+^ expression was greater in male mice when compared to female mice (**Fig. 3P**). AMG c-Fos^+^ activation was similar among the groups except in the COA and PAA (**Fig. 3Q**). These data suggest that the HPC and AMG are impacted by sex and AD progression.

### Female AD mice exhibit a decrease in memory trace activation in dorsal CA3

c-Fos^+^ is a proxy of neural activity and provides a single timepoint of activity in relation to behavior. To better understand how female and male AD mice differ in their encoding and retrieving of individual memories, APP/PS1 mice were bred with the ArcCreER^T2^ x ChR2-eYFP model, to allow for indelible activity-dependent tagging. Thus, multiple time points of activity could be identified (**Fig. 4A, Table S3**). Mice were injected with 4-OHT 5 hours before 3-shock CFC to indelibly tag active ensembles. Five days later, mice were re-exposed to the same context and euthanized 1 hour later (**Fig. 4B**). Both female and male AD mice exhibited cognitive decline at 6 months of age compared to Ctrl mice (**Fig. 4C-D**). Plaques were stained with Methoxy-X04 to confirm AD pathology (**Fig. 4E-F**) and memory traces were present throughout the HPC (**Fig. 4G-4I)**.

**Fig 4.**
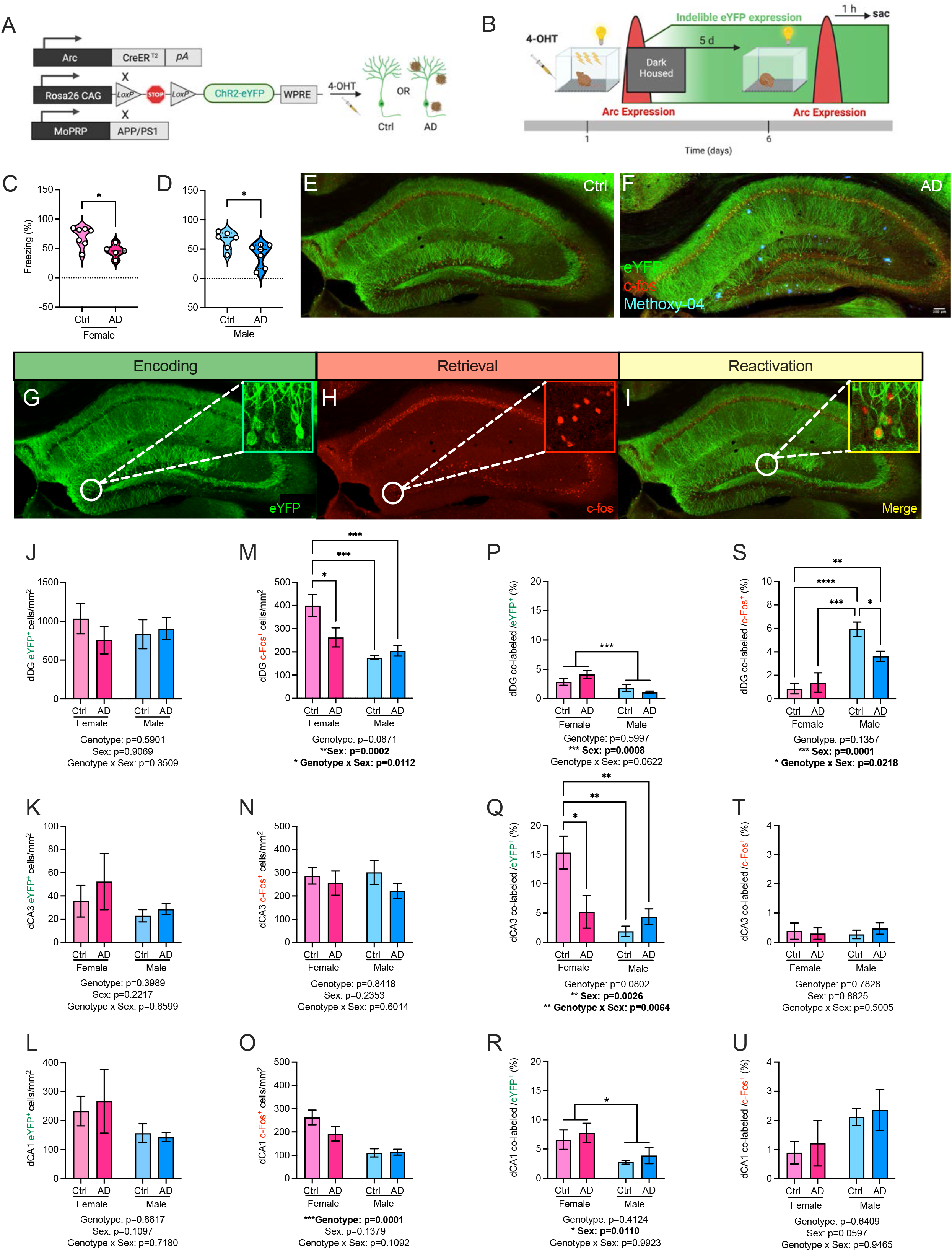
Female AD mice exhibit a decrease in memory traces in the dorsal CA3. (**A**) Genetic design. (**B**) Experimental timeline. (**C-D**) Female and male AD mice exhibit decreased freezing behavior at 6 months of age compared to Ctrl mice. (**E-F**) Representative images confirming plaque pathology in AD mice using Methoxy-X04. (**G-I**) Representative images of indelible memory tagging in the dHPC. (**J-L**) Female Ctrl and AD mice exhibit comparable levels of eYFP^+^-tagged cells in the dDG, dCA3, and dCA1. (**M**) In the dDG, c-Fos^+^ activation is decreased in female AD mice, and Ctrl and AD male mice. (**N-O**) c-Fos^+^ activation is similar among the groups in dCA3 and dCA1. (**P**) In the dDG, female mice exhibit a greater co-labeled / eYFP (%) when compared to male AD mice. (**Q**) In dCA3, female AD, and Ctrl and AD male mice a decreased co-labeled / eYFP (%) when compared to Ctrl female mice. (**R**) In dCA1, female mice exhibit a greater co-labeled / eYFP (%) when compared to male mice. (**S**) in dDG, there are no differences in female groups for co-labeled / c-Fos^+^ (%). Male AD mice exhibit a decreased co-labeled / c-Fos^+^ (%) when compared to male Ctrl mice. (**T-U**) Memory trace activation is similar among the groups in dCA3 and dCA1. (n=4-7 mice per group). Error bars represent ± SEM. *p<0.05, **p<0.01, ***p<0.001. pA, polyadenylation; ChR2, channelrhodopsin-2; eYFP, enhance yellow fluorescent protein; WPRE, woodchuck hepatitis virus post-transcriptional regulatory element; 4-OHT, 4-hydroxytamoxifen; d, days; h, hours; sac, sacrifice; Ctrl, control; AD, Alzheimer’s disease; mm, millimeter; dDG, dorsal dentate gyrus; dCA3, dorsal cornu ammonis; dCA1, dorsal cornu ammonis; CA1.

For eYFP^+^, all groups showed comparable levels in the dDG, dCA3, and dCA1 (**Fig. 4J-4L**). In the dDG, female AD mice exhibited decreased c-Fos^+^ expression; male mice exhibited less c-Fos^+^ expression than female mice, similar to the network trends (**Fig. 4M**). c-Fos^+^ expression was similar between groups in dCA3 and dCA1 (**Fig. 4N-O**). In the dDG, dCA3, and dCA1, there was an effect of Sex on the co-labeled/eYFP^+^ (%) (**Fig. 4P-4R**). However, in dCA3, female AD mice exhibited decreased reactivation (**Fig. 4Q**). In the dDG, male mice exhibited a greater co-labeled/c-Fos^+^ (%) compared to female mice; male AD male exhibited less reactivation compared to Ctrl mice (**Fig. 4S**). These results corroborate with our previous findings in male AD mice (13). All groups had comparable co-labeled/c-Fos^+^ (%) in dCA3 and dCA1 (**Fig. 4T-4U**). In summary, AD impacts memory traces throughout the dorsal HPC in a sex-specific manner.

### c-Fos^+^ activation is increased in the vCA1of AD female mice during memory retrieval

Memory traces were next quantified in the vHPC, as this region is important for fear memory and anxiety-related behaviors (**Fig. 5A-F)**. In the vDG and vCA3, eYFP^+^ and c-Fos^+^ expression was similar between all the groups (**Fig. 5G-5H**, **Fig. 5J-5K**). In vCA1, female AD female mice: 1) exhibited increased reactivation compared to male mice (**Fig. 5I**) and 2) had increased c-Fos^+^ expression compared to Ctrl mice (**Fig. 5L**). The percentage of co-labeled/eYFP^+^ cells was comparable in all groups in all regions (**Fig. 5M-O**). However, there was a significant effect of Sex on co-labeled/c-Fos^+^ (%) in vDG and vCA1 (**Fig. 5P, 5R**), but not in vCA3 (**Fig. 5Q**), with male mice having increased reactivation relative to female mice. These data suggest that dHPC memory traces are impacted more by AD than vHPC memory traces.

**Fig 5.**
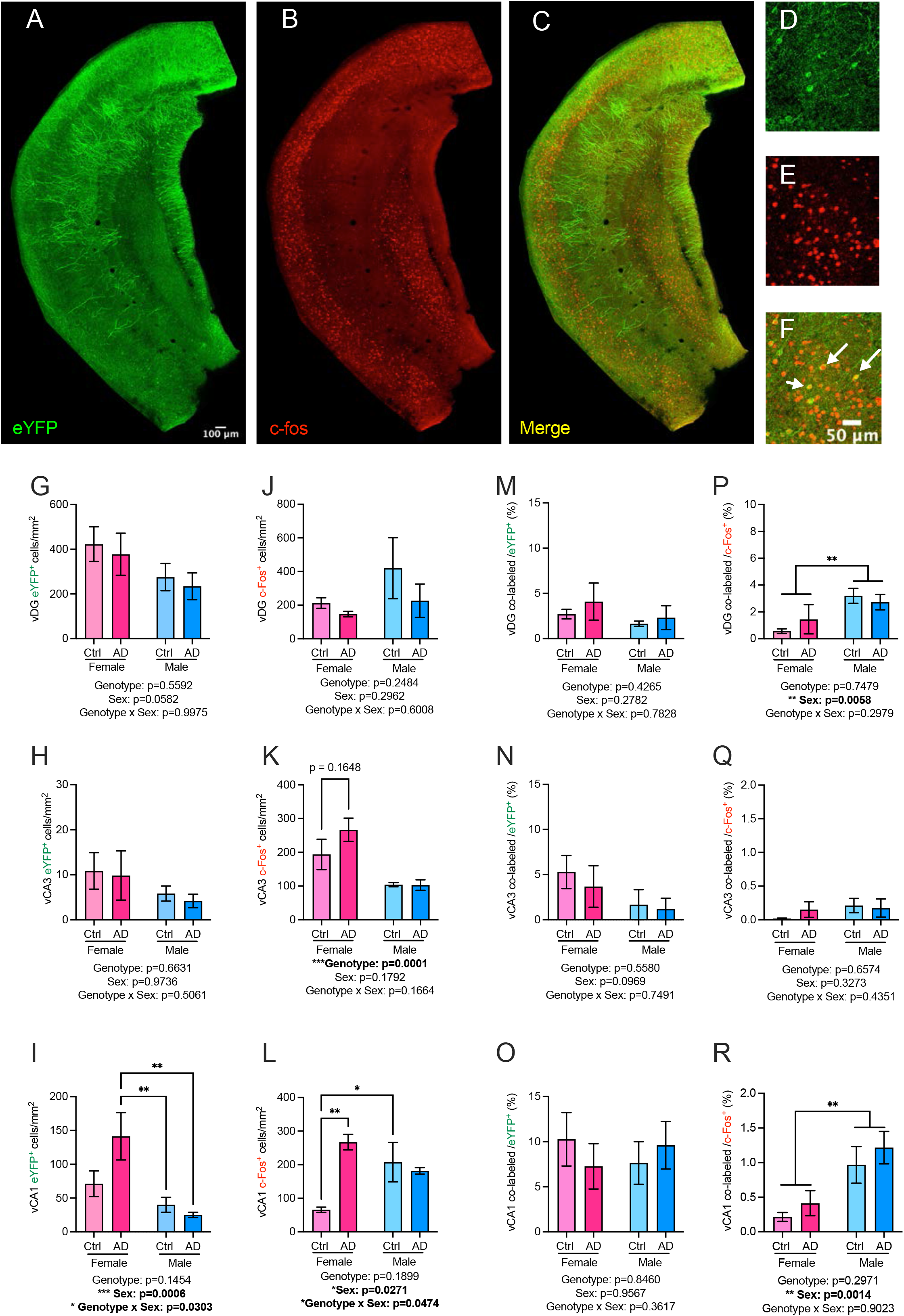
Encoding and retrieval cell activation are increased in the ventral hippocampus of AD female mice. (**A-F**) Representative images of eYFP^+^, c-Fos^+^, and memory traces in the vHPC. (**G-H**) eYFP expression was comparable in the groups in the vDG and vCA3. (**I**) In vCA1, female AD mice exhibit increased eYFP^+^ activation compared to male Ctrl and AD mice. (**J-K**) c-Fos^+^ expression was comparable in the groups in the vDG and vCA3. (**L**) In vCA1, female AD mice exhibited increased c-Fos^+^ expression compared to female Ctrl mice. Male Ctrl mice exhibited increased c-Fos^+^ activation when compared to female Ctrl mice. (**M-O**) In vDG, vCA3, and vCA1, the co-labeled / eYFP (%) was comparable between all groups. (**P**) Male mice exhibited a greater co-labeled / c-Fos^+^ (%) when compared to female mice. (**Q**) In vCA3, the co-labeled / c-Fos^+^ (%) was comparable between all groups. (**R**) In vCA1, male mice exhibited a greater co-labeled / c-Fos^+^ (%) than female mice. (n=4-7 mice per group). Error bars represent ± SEM. *p<0.05, **p<0.01, ***p<0.001. eYFP, enhanced yellow fluorescent protein; mm, microns; mm, millimeter; vDG, ventral dentate gyrus; vCA3, ventral cornu ammonis 3; vCA1, ventral cornu ammonis 1; Ctrl, control; AD, Alzheimer’s disease.

### Female patients with anxiety and amyloid transition more quickly to dementia

To determine whether our results could be translated to the human population, we analyzed data from the ADNI cohort. A total 1661 participants were included with baseline Neuropsychiatric Inventory Questionnaire (NPIQ) anxiety subscales (**Fig. 6A, Table S4**). In both men and women, mild cognitive impairment (MCI) and AD participants showed higher anxiety prevalence compared to cognitively normal (CN) participants (**Table S5**). Men showed a greater increase from CN to AD, but this was mainly due to their low baseline anxiety (**Fig. 6B-6C**). Additionally, Age was significant, but the interaction between Sex and Diagnosis group was at trend level. The odds ratio of having anxiety was 8x and 24x greater in female and male AD participants compared to CN, respectively (**Fig. 6D and Table S6**). These data indicate that women and men with anxiety are at increased risk for developing dementia.

**Fig. 6.**
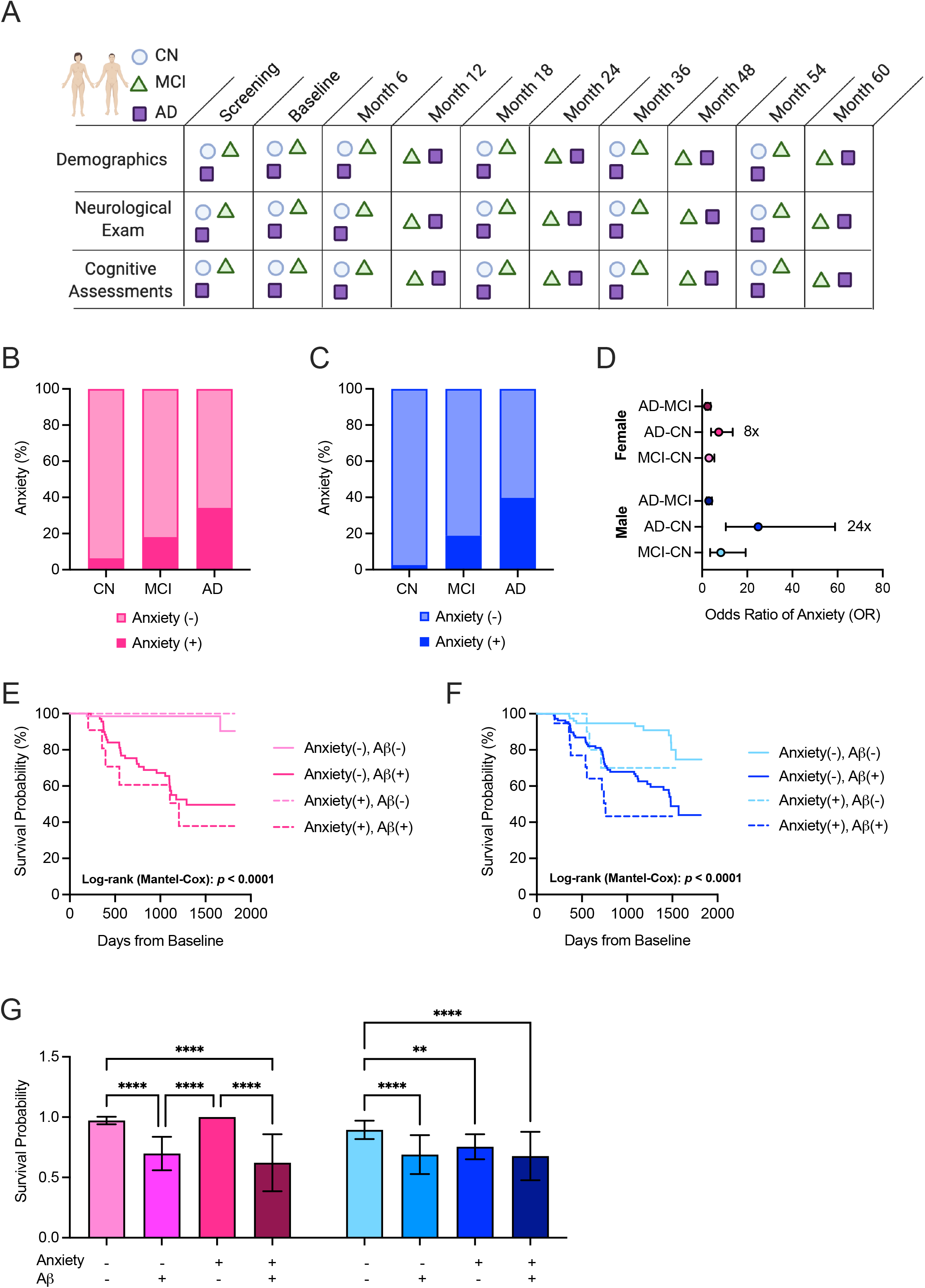
Female patients with anxiety and amyloid transition more quickly to dementia. **(A)** ADNI data collection pipeline. **(B-C)** The percentage of subjects with anxiety is increased in those with MCI and AD in both females and males. **(D)** The odds ratio indicates that female AD subjects are eight times more likely to have anxiety compared to CN subjects, and male AD subjects are 24 times more likely to have anxiety compared to CN subjects. (**E-F**) Female subjects with amyloid and anxiety transition faster to dementia than males. (**G**) Bar graph representation of the survival plots in **E-F** indicates that subjects with amyloid positivity and anxiety transition to dementia at a faster rate. (n=1661). Error bars represent ± SEM. *p<0.05, **p<0.01, ***p<0.001. CN, cognitively normal; MCI, mild cognitive impairment; AD, Alzheimer’s disease; OR, odds ratio of anxiety; Aβ, amyloid beta.

Among 823 MCI participants at baseline, 418 were followed with CSF measures. For consistency across subjects followed in the ADNI study, we limited up-to a 5 year follow up. Among 418 MCI participants, 113 (27.36%) subjects transitioned to AD in 5 years. Having anxiety and controlling for age, gender, and education, we further explored the gender interactions with anxiety and A*β* positivity. A*β* positivity had significantly higher hazard ratio (HR) in females than males (**Fig. 6E-6G, Table S7 and S8**). There was no three-way interaction. These data indicate that women with anxiety are at increased risk for amyloid accumulation and dementia transition.

### Women with anxiety exhibit decreased amygdala and hippocampal volume

Next, we tested whether brain volumes differed by anxiety group. Subjects with anxiety showed large volumes in left pars trangularis (inferior frontal gyrus), bilateral pericalcarine (visual cortex), right choroid plexus (ventricles), and right pallidum, while smaller corpus callosum (CC)-central and CC-posterior. However, the group difference did not survive multiple comparison correction. Sex did influence brain volumes (**Table S9**). fMRI images between male and female subjects anatomically show differences in brain volume (**Fig. 7A**). Using Elasticnet, we evaluated if anxiety and brain atrophy could predict the transition to dementia. Based on repeated 5-fold cross-validation in 10 random initializations of cross-validation, there were 23 ROIs that were selected at least once in the model, and their average importance scores and average coefficients are reported in **Table S10**. Anxiety was the most important variable to predict dementia transition above brain atrophy and age (**Fig. 7B**). Using sparse group lasso, we evaluated brain volume and anxiety interactions as well as sex by brain volume interaction. Interestingly, females with anxiety showed smaller volumes while males tended to have larger brain volumes in the right inferior temporal gyrus and AMG (**Fig. 7C-7D**). In the right insula and bilateral ventral diencephalon, only males with anxiety showed increased volume (**Fig. 7E-7G**). Decreased volume was observed in the right HPC in females (**Fig. 7H**). We also tested diagnosis by anxiety and the three-way interaction of anxiety, diagnosis, and sex but the regions did not survive multiple comparison correction. These data indicate anxiety as the best predictor of dementia transition above brain atrophy.

**Fig. 7.**
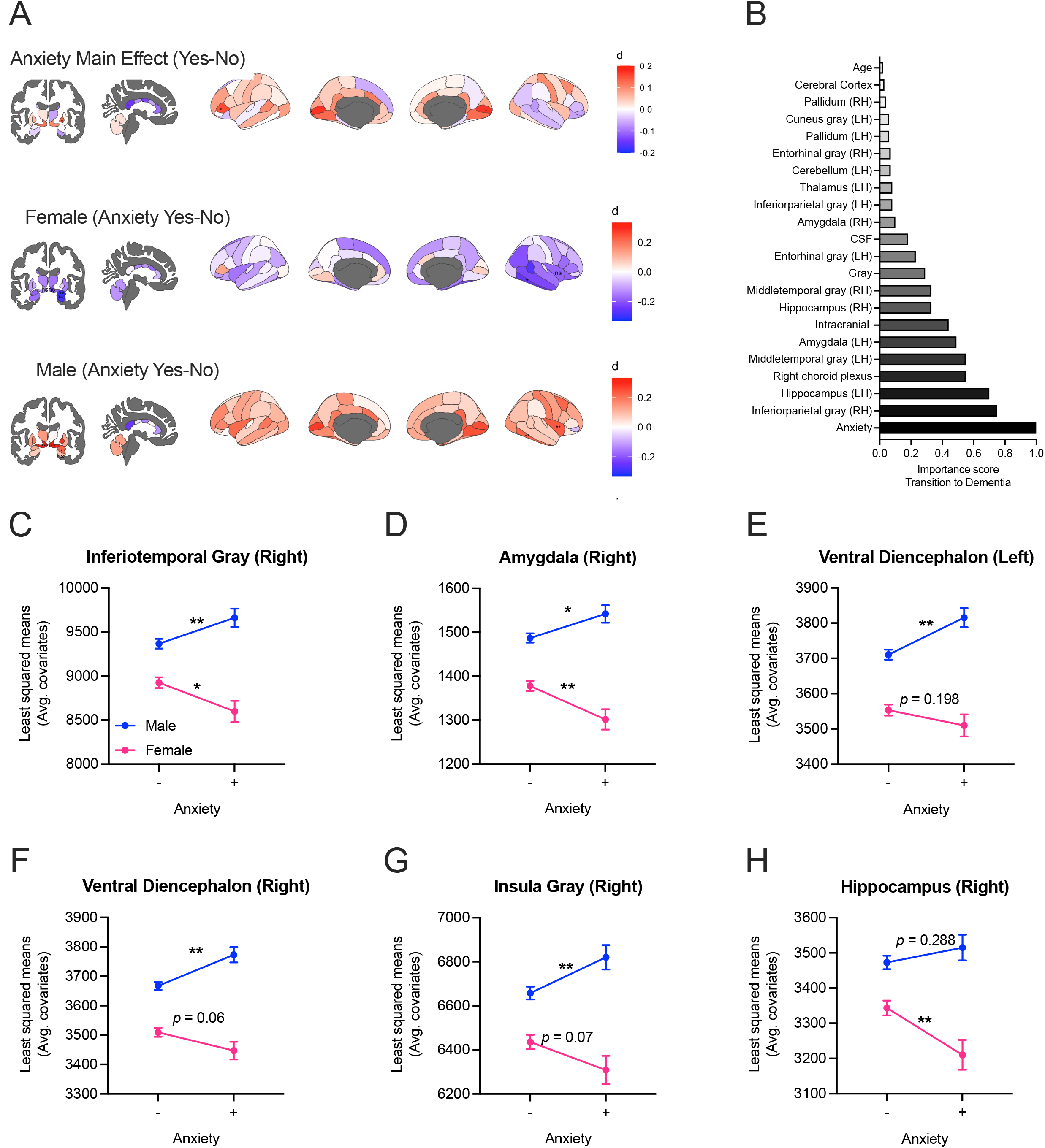
Sex-specific effect of anxiety on brain volume in ADNI participants. (**A**) fMRI representative images of female-male, female, and male brain volumes using a t-value scale (red= increased volume, blue=decreased volume). **(B)** Anxiety is the greatest predictor of dementia transition in male and female subjects compared to brain atrophy and age. **(C-D)** Female subjects with anxiety have decreased inferior temporal gray volume and amygdala volume compared to female subjects without anxiety. Male subjects with anxiety have increased inferior temporal gray and amygdala volume compared to male subjects without anxiety. (**E**) Male subjects with anxiety have increased ventral diencephalon volume in the left hemisphere compared to male subjects without anxiety. There is no significant difference in the female groups. (**F**) Male subjects with anxiety have increased ventral diencephalon volume in the right hemisphere compared to male subjects without anxiety. There is no significant difference in the female groups, although it is trending. (**G**) In the right hemisphere, male subjects with anxiety have increased insula gray volume. There is no significant difference in the female groups, although it is trending. (**H**) Female subjects with anxiety have decreased hippocampal volume. There is no difference between the groups for male subjects in the hippocampus. (n=1650). Error bars represent ± SEM. *p<0.05, **p<0.01, ***p<0.001.

## DISCUSSION

Here, we explored the relationship between anxiety and sex, and the behavioral and biological impact on AD progression. While it is well known that cognition is greatly impacted by AD, the role of anxiety in AD progression is less defined (13,17,18,28–30). We found that: 1) female AD mice exhibited anxiety-like behavior and cognitive decline at an earlier age than Ctrl and male mice, 2) network connectivity is impacted in a sex-specific manner in AD mice, 3) sex differentially impacts memory traces in the HPC of AD mice, 4) anxiety significantly influences the transition to dementia in human subjects, and 5) female subjects positive for anxiety and amyloid transitioned more quickly to dementia than male subjects.

We report that female AD mice exhibit increased anxiety-like behavior. However, these data are in contrast with a recent study using the 3xTg-AD mice, which showed decreased anxiety. This difference is most likely to do the lack of female mice and to alterations in in locomotion in the 3xTg-AD mouse line (28). In APP/PS1 mice, others have shown that 4- and 24-month-old male mice exhibit increased anxiety in the OF and EPM (12,16). The lack of anxiety at younger ages in our AD male mice could be due to strain differences between the studies. Since the APP/PS1 mice display NPS in both sexes and at an early age, this model may be better suited for studying the psychiatric symptoms of patients with early-stage AD. Further research will be necessary to determine whether this anxiety-like behavior can be rescued with novel anxiolytics or cognitive-enhancing compounds.

Female AD mice exhibited impaired cognition at an earlier age (i.e., at 2 months of age) than male AD mice (i.e., at 6 months of age) The earlier cognitive decline in AD female mice is common and often reported in both the APP/PS1 and 3xTg-AD models where females displayed greater plaque burden (31) and earlier memory deficits in an alternation task (32,33). However, when males were demasculinized and females were defeminized, the opposite effects were observed, suggesting a role for sex-steroid hormones in AD progression (33). Future studies will determine whether hormone replacement therapy (HRT) can buffer not only the cognitive, but psychiatric phenotypes observed in the APP/PS1 mice.

Brain-wide analysis of c-Fos^+^ revealed changes in regional correlations and overall network connectivity in AD mice. Because this is the first study to analyze brain-wide IEG activity in male and female AD mice, we compared our results to those using fMRI and functional connectivity patterns. fMRI studies examining female Tau models of AD show decreased functional state connectivity (34), similar to our results showing sparse connections in female AD mice. Additionally, decreased interhemispheric hippocampal connectivity has been reported in female 3xTg-AD mice (35). In Ctrl mice, female mice displayed increased connectivity compared to male mice, which is in agreement with a recent study reporting increased connectivity in female mice with age (36). Of particular interest, male AD mice displayed opposite connectivity patterns compared to female AD mice, possibly due to hyperexcitability, often observed in early-stage AD (24). Others note that in male mice, APOE4, the dominant risk allele for AD, enhances neuronal hyperactivity due to a loss of inhibitory tone in the ECT (37). While female AD mice may still experience hyperexcitability, it may occur at an earlier age (i.e., when anxiety is increased). In agreement with this hypothesis, a recent study reported increased connectivity followed by a later decrease in female APP knock-in mice (36). Future studies will examine earlier timepoints for brain-wide connectivity patterns.

We replicated our previous findings that memory traces are decreased in the dDG of male AD mice (7), but now additionally report that memory traces are decreased in dCA3 of female AD mice. While we and others have shown that optogenetic stimulation of the dDG rescues memory in AD mice (7,38,39), there has been lack of inclusion of female AD mice in these studies. In addition to the DG, we also report sex-specific differences in vCA1 with increased eYFP^+^ and c-Fos^+^ expression in female AD mice. vCA1 is enriched with cells that are activated by anxiogenic environments and required for avoidance behavior (40), and the observed increase in activity that we report may also contribute to the female-specific increase in anxiety-like behavior. Future studies will focus on elucidating ventral hippocampal abnormalities in a sex-specific manner.

AD subjects displayed higher levels of anxiety compared to CN and MCI subjects, similar to our rodent findings. These results coincide with other studies showing that depression and anxiety are commonly observed features of MCI which are linked to cognitive and functional decline (41,42). A recent Swedish BioFinder study assessed anxiety and apathy as early markers of AD and found that both were related to amyloid deposition and predicted cognitive decline (43). However, there was no analysis of sex differences. Here, we provide additional evidence that women with CSF amyloid and anxiety symptoms transition at a faster rate to dementia compared to men. Although our analysis was limited to amyloid, others have proven this for tau pathology as well. Women with AD have significantly higher tau pathology across multiple regions of the cortical mantle, which contributes to faster cognitive decline (44). Therefore, examining how anxiety is related to tau pathology is a critical future direction.

Volumetric changes in the brain have been controversial with reports of decreased amygdala and hippocampal volumes in depressed patients (45), and others finding increased volumes with little report on sex differences (46). The differences we observed in brain volume might not be a significant contributor to the functional behavior outcomes. How the network and functional connectivity change rather than volumetric changes will most likely be more important when determining the contribution of these brain regions in anxiety and AD progression (47).

In summary, these findings elucidate the role of anxiety and sex in AD progression in mice and humans. While future studies are needed to understand whether anxiety is a predictor, a NPS biomarker, or a comorbid symptom that occurs during disease onset, it is evident that anxiety impacts and exacerbates the transition to dementia. These results suggest that AD network dysfunction is sexually dimorphic, and that personalized medicine may benefit male and female AD patients rather than a one size fits all approach.

## ACKNOWLEDGMENTS AND DISCLOSURES

The research reported in the article was supported by grants from the NIMH (Grant No. 2T32 MH015174-40 [to CAD and HCH]), the NIA (Grant No. K99 AG059953-01A1 [to HCH], Grant No. R24 AG061421 [to HCH], Grant No. R21 AG064774-02 [to CAD], and Grant No. R56 AG058661-01 [to CAD]), the NIH (Grant No. DP5 OD017908-01 [to CAD]), and the NICHD (Grant No. R01 HD101402-01 [to CAD]). This work was also funded by a Faculty Research Fellowship Relating to the Study of Optimal Aging from the Columbia Aging Center at Columbia University Irving Medical Center (CUIMC) [to CAD].

Data collection and sharing for this project was funded by the Alzheimer’s Disease Neuroimaging Initiative (ADNI) (National Institutes of Health Grant U01 AG024904) and DOD ADNI (Department of Defense award number W81XWH-12-2-0012). ADNI is funded by the National Institute on Aging, the National Institute of Biomedical Imaging and Bioengineering, and through generous contributions from the following: AbbVie, Alzheimer’s Association; Alzheimer’s Drug Discovery Foundation; Araclon Biotech; BioClinica, Inc.; Biogen; Bristol-Myers Squibb Company; CereSpir, Inc.; Cogstate; Eisai Inc.; Elan Pharmaceuticals, Inc.; Eli Lilly and Company; EuroImmun; F. Hoffmann-La Roche Ltd and its affiliated company Genentech, Inc.; Fujirebio; GE Healthcare; IXICO Ltd.; Janssen Alzheimer Immunotherapy Research & Development, LLC.; Johnson & Johnson Pharmaceutical Research & Development LLC.; Lumosity; Lundbeck; Merck & Co., Inc.; Meso Scale Diagnostics, LLC.; NeuroRx Research; Neurotrack Technologies; Novartis Pharmaceuticals Corporation; Pfizer Inc.; Piramal Imaging; Servier; Takeda Pharmaceutical Company; and Transition Therapeutics. The Canadian Institutes of Health Research is providing funds to support ADNI clinical sites in Canada. Private sector contributions are facilitated by the Foundation for the National Institutes of Health (www.fnih.org). The grantee organization is the Northern California Institute for Research and Education, and the study is coordinated by the Alzheimer’s Therapeutic Research Institute at the University of Southern California. ADNI data are disseminated by the Laboratory for Neuro Imaging at the University of Southern California.

We thank the members of the Denny laboratory, the biostatistics team, and Dr. Haley Shelley for their insightful comments on this project and manuscript.

CAD is named on a provisional patent application for the prophylactic use of (*R*,*S*)-ketamine and other compounds against stress-related psychiatric disorders. All other authors report no biomedical financial interests or potential conflicts of interest.

## AUTHOR CONTRIBUTIONS

HCH was responsible for the experimental idea, design, performing experiments, data analysis, figures, and manuscript writing. SL was responsible for the ADNI dataset analysis and figures and writing of the methods and results. ML and MJ created the whole-brain pipeline and helped with analyzing cell counts. MJ edited the pipeline and added new clustering graphs. JC was part of the ADNI dataset team. MS and AW ran analysis for the cell counts. KJ ran analysis for cell counts and helped to write the behavioral results section. CAD mentored designed the experiments, analyzed data, created figures, and edited the manuscript.

## REFERENCES

1. 2021 Alzheimer’s disease facts and figures (2021): Alzheimers Dement 17: 327–406.

2. Beason-Held LL, Goh JO, An Y, Kraut MA, O’Brien RJ, Ferrucci L, Resnick SM (2013): Changes in Brain Function Occur Years before the Onset of Cognitive Impairment. J Neurosci 33: 18008–18014.

3. Mah L, Binns MA, Steffens DC, Alzheimer’s Disease T, Initiative N (2014): Anxiety symptoms in amnestic mild cognitive impairment are associated with medial temporal atrophy and predict conversion to Alzheimer’s disease. Am J Geriatr Psychiatry 23: 466–476.

4. Crimmins EM, Shim H, Zhang YS, Kim JK (2019): Differences between Men and Women in Mortality and the Health Dimensions of the Morbidity Process. Clin Chem 65: 135.

5. Callahan MJ, Lipinski WJ, Bian F, Durham RA, Pack A, Walker LC (2001): Augmented Senile Plaque Load in Aged Female β-Amyloid Precursor Protein-Transgenic Mice. Am J Pathol 158: 1173–1177.

6. Denny CA, Kheirbek MA, Alba EL, Tanaka KF, Brachman RA, Laughman KB, et al. (2014): Hippocampal memory traces are differentially modulated by experience, time, and adult neurogenesis. Neuron 83: 189–201.

7. Perusini JN, Cajigas SA, Cohensedgh O, Lim SC, Pavlova IP, Donaldson ZR, Denny CA (2017): Optogenetic stimulation of dentate gyrus engrams restores memory in Alzheimer’s disease mice. Hippocampus 27: 1110–1122.

8. Madisen L, Zwingman TA, Sunkin SM, Oh SW, Zariwala HA, Gu H, et al. (2010): A robust and high-throughput Cre reporting and characterization system for the whole mouse brain. Nat Neurosci 13: 133–140.

9. Denny CA, Kheirbek MA, Alba EL, Tanaka KF, Brachman RA, Laughman KB, et al. (2014): Hippocampal memory traces are differentially modulated by experience, time, and adult neurogenesis. Neuron 83: 189–201.

10. Weiner MW, Veitch DP, Aisen PS, Beckett LA, Cairns NJ, Green RC, et al. (2013): The Alzheimer’s Disease Neuroimaging Initiative: a review of papers published since its inception. Alzheimers Dement 9: e111–e194.

11. Pugh PL, Richardson JC, Bate ST, Upton N, Sunter D (2007): Non-cognitive behaviours in an APP/PS1 transgenic model of Alzheimer’s disease. Behav Brain Res 178: 18–28.

12. Huang H, Nie S, Cao M, Marshall C, Gao J, Xiao N, et al. (2016): Characterization of AD-like phenotype in aged APPSwe/PS1dE9 mice. Age 38: 303–322.

13. Arendash GW, King DL, Gordon MN, Morgan D, Hatcher JM, Hope CE, Diamond DM (2001): Progressive, age-related behavioral impairments in transgenic mice carrying both mutant amyloid precursor protein and presenilin-1 transgenes. Brain Res 891: 42–53.

14. Kosel F, Pelley JMS, Franklin TB (2020): Behavioural and psychological symptoms of dementia in mouse models of Alzheimer’s disease-related pathology. Neurosci Biobehav Rev 112: 634–647.

15. Holcomb L, Gordon MN, Mcgowan E, Yu X, Benkovic S, Jantzen P, et al. (1998): Accelerated Alzheimer-type phenotype in transgenic mice carrying both mutant amyloid precursor protein and presenilin 1 transgenes. Nat Med 4: 97–100.

16. Gao JY, Chen Y, Su DY, Marshall C, Xiao M (2018): Depressive- and anxiety-like phenotypes in young adult APPSwe/PS1dE9 transgenic mice with insensitivity to chronic mild stress. Behav Brain Res 353: 114–123.

17. Bedrosian TA, Herring KL, Weil ZM, Nelson RJ (2011): Altered temporal patterns of anxiety in aged and amyloid precursor protein (APP) transgenic mice. Proc Natl Acad Sci U S A 108: 11686–11691.

18. Bruce-Keller AJ, Gupta S, Knight AG, Beckett TL, McMullen JM, Davis PR, et al. (2011): Cognitive impairment in humanized APP×PS1 mice is linked to Aβ(1-42) and NOX activation. Neurobiol Dis 44: 317–326.

19. Thomas A, Burant A, Bui N, Graham D, Yuva-Paylor LA, Paylor R (2009): Marble burying reflects a repetitive and perseverative behavior more than novelty-induced anxiety. Psychopharmacology (Berl) 204: 361.

20. Pavlova IP, Shipley SC, Lanio M, Hen R, Denny CA (2018): Optimization of immunolabeling and clearing techniques for indelibly-labeled memory traces. Hippocampus 28: 523–535.

21. Leal Santos S, Stackmann M, Muñoz Zamora A, Mastrodonato A, De Landri A V., Vaughan N, et al. (2021): Propranolol decreases fear expression by modulating fear memory traces. Biol Psychiatry 89: 1150–1161.

22. Kim W Bin, Cho JH (2017): Synaptic Targeting of Double-Projecting Ventral CA1 Hippocampal Neurons to the Medial Prefrontal Cortex and Basal Amygdala. J Neurosci 37: 4868–4882.

23. Xu C, Krabbe S, Gründemann J, Botta P, Fadok JP, Osakada F, et al. (2016): Distinct Hippocampal Pathways Mediate Dissociable Roles of Context in Memory Retrieval. Cell 167: 961–972.e16.

24. Reagh ZM, Noche JA, Tustison NJ, Delisle D, Murray EA, Yassa MA (2018): Functional Imbalance of Anterolateral Entorhinal Cortex and Hippocampal Dentate/CA3 Underlies Age-Related Object Pattern Separation Deficits. Neuron 97: 1187–1198.e4.

25. Poirier GL, Amin E, Good MA, Aggleton JP (2011): Early-onset dysfunction of retrosplenial cortex precedes overt amyloid plaque formation in Tg2576 mice. Neuroscience 174: 71–83.

26. Sosulina L, Mittag M, Geis HR, Hoffmann K, Klyubin I, Qi Y, et al. (2021): Hippocampal hyperactivity in a rat model of Alzheimer’s disease. J Neurochem 157: 2128–2144.

27. Perusini JN, Cajigas SA, Cohensedgh O, Lim SC, Pavlova IP, Donaldson ZR, Denny CA (2017): Optogenetic stimulation of dentate gyrus engrams restores memory in Alzheimer’s disease mice. Hippocampus 27: 1110–1122.

28. Várkonyi D, Török B, Sipos E, Fazekas CL, Bánrévi K, Correia P, et al. (2022): Investigation of Anxiety- and Depressive-like Symptoms in 4- and 8-Month-Old Male Triple Transgenic Mouse Models of Alzheimer’s Disease. Int J Mol Sci 23: 10816.

29. Tag SH, Kim B, Bae J, Chang KA, Im HI (2022): Neuropathological and behavioral features of an APP/PS1/MAPT (6xTg) transgenic model of Alzheimer’s disease. Mol Brain 15: 1–13.

30. Yang JT, Wang ZJ, Cai HY, Yuan L, Hu MM, Wu MN, Qi JS (2018): Sex Differences in Neuropathology and Cognitive Behavior in APP/PS1/tau Triple-Transgenic Mouse Model of Alzheimer’s Disease. Neurosci Bull 34: 736–746.

31. Wang J, Tanila H, Puoliväli J, Kadish I, Van Groen T (2003): Gender differences in the amount and deposition of amyloidbeta in APPswe and PS1 double transgenic mice. Neurobiol Dis 14: 318–327.

32. Arsenault D, Tremblay C, Emond V, Calon F (2020): Sex-dependent alterations in the physiology of entorhinal cortex neurons in old heterozygous 3xTg-AD mice. Biol Sex Differ 11: 1–19.

33. Carroll JC, Rosario ER, Kreimer S, Villamagna A, Gentzschein E, Stanczyk FZ, Pike CJ (2010): Sex differences in β-amyloid accumulation in 3xTg-AD mice: Role of neonatal sex steroid hormone exposure. Brain Res 1366: 233.

34. Green C, Sydow A, Vogel S, Anglada-Huguet M, Wiedermann D, Mandelkow E, et al. (2019): Functional networks are impaired by elevated tau-protein but reversible in a regulatable Alzheimer’s disease mouse model. Mol Neurodegener 14: 1–13.

35. Manno FAM, Isla AG, Manno SHC, Ahmed I, Cheng SH, Barrios FA, Lau C (2019): Early Stage Alterations in White Matter and Decreased Functional Interhemispheric Hippocampal Connectivity in the 3xTg Mouse Model of Alzheimer’s Disease. Front Aging Neurosci 11: 39.

36. Morrissey ZD, Gao J, Zhan L, Li W, Fortel I, Saido T, et al. (2023): Hippocampal functional connectivity across age in an App knock-in mouse model of Alzheimer’s disease. Front Aging Neurosci 14: 1085989.

37. Nuriel T, Angulo SL, Khan U, Ashok A, Chen Q, Figueroa HY, et al. (2017): Neuronal hyperactivity due to loss of inhibitory tone in APOE4 mice lacking Alzheimer’s disease-like pathology. Nat Commun 8: 1464.

38. Etter G, van der Veldt S, Manseau F, Zarrinkoub I, Trillaud-Doppia E, Williams S (2019): Optogenetic gamma stimulation rescues memory impairments in an Alzheimer’s disease mouse model. Nat Commun 10: 5322.

39. Roy DS, Arons A, Mitchell TI, Pignatelli M, Ryan TJ, Tonegawa S (2016): Memory retrieval by activating engram cells in mouse models of early Alzheimer’s disease. Nature 531: 508– 512.

40. Jimenez JC, Su K, Goldberg AR, Luna VM, Biane JS, Ordek G, et al. (2018): Anxiety Cells in a Hippocampal-Hypothalamic Circuit. Neuron 97: 670–683.

41. Mah L, Binns MA, Steffens DC (2015): Anxiety Symptoms in Amnestic Mild Cognitive Impairment Are Associated with Medial Temporal Atrophy and Predict Conversion to Alzheimer Disease. Am J Geriatr Psychiatry 23: 466–476.

42. Ma L (2020): Depression, Anxiety, and Apathy in Mild Cognitive Impairment: Current Perspectives. Front Aging Neurosci 12.

43. Johansson M, Stomrud E, Lindberg O, Westman E, Johansson PM, van Westen D, et al. (2020): Apathy and anxiety are early markers of Alzheimer’s disease. Neurobiol Aging 85: 74–82.

44. Buckley RF, Scott MR, Jacobs HIL, Schultz AP, Properzi MJ, Amariglio RE, et al. (2020): Sex Mediates Relationships Between Regional Tau Pathology and Cognitive Decline. Ann Neurol 88: 921–932.

45. Lange C, Irle E (2004): Enlarged amygdala volume and reduced hippocampal volume in young women with major depression. Psychol Med 34: 1059–1064.

46. MacMillan S, Szeszko PR, Moore GJ, Madden R, Lorch E, Ivey J, et al. (2003): Increased amygdala: hippocampal volume ratios associated with severity of anxiety in pediatric major depression. J Child Adolesc Psychopharmacol 13: 65–73.

47. Gold AL, Shechner T, Farber MJ, Spiro CN, Leibenluft E, Pine DS, Britton JC (2016): Amygdala-cortical connectivity: Associations with anxiety, development, and threat. Depress Anxiety 33: 917–926.

