## Supplement for "Sex-Specific Effects of Anxiety on Cognition and Activity-Dependent Neural Networks: Insights from (Female) Mice and (Wo)Men"

### SUPPLEMENTARY METHODS

### MOUSE METHODS

#### Behavioral Tests

Behavioral testing occurred during the light phase and mice underwent all tests at 2, 4, or 6 months of age. All experimenters were blind to the genotype of individual mice.

***Open Field (OF)***

The OF assay was administered as previously described to assess general motor activity (1,2). Briefly, mice were placed into an open arena, 50 cm x 50 cm x 50 cm, and allowed to explore for 10 minutes. Distance and time spent in the center and periphery were measured and using MotorMonitor software (Kinder Scientific, Poway, CA) as an indicator of anxiety-like behavior. The floor and walls were cleaned with 70% EtOH and dried after each session. Age and Genotype were analyzed using a two-way ANOVA with Tukey post-hoc tests.

***Elevated Plus Maze (EPM)***

The EPM is a widely used behavioral assay for rodents to assess the anxiolytic effects of pharmacological agents and to define brain regions and mechanisms underlying anxiety-related behavior (3). The EPM was administered as previously described (1,2). Briefly, the EPM was conducted in a plus-cross-shaped apparatus that was 50 cm above the ground. Two arms of the maze are enclosed with walls while 2 arms are open. Mice were placed in the center of the apparatus facing an open arm and allowed to explore for 5 minutes. ANY-maze (Stoelting Co., Wood Dale, IL) software was used to track the amount of time spent in the open arms versus the closed arms. Age and Genotype were analyzed using a two-way ANOVA with Tukey post-hoc tests.

***Marble Burying (MB)***

The MB assay was conducted in a clean cage (10.5 in x 5.5 in) containing soft pliable Beta Chip bedding (Northeastern Products Corp, Warrensburg, NY). The cage contained 16 marbles set up in 4 rows of 4 across. Mice were given 30 minutes to explore and bury. At the end of the assay, the percentage of marbles buried was calculated. Age and Genotype were analyzed using a two-way ANOVA with Tukey post-hoc tests.

***Light Dark Test (LDT)***

Mice were placed on the light side of a shuttle box avoidance chamber (Med Associates, Fairfax, VT) and after 5 seconds a door opened and mice were allowed to explore the dark side. The animals could move freely between the light and dark side for 10 minutes. The number of beam breaks was counted as activity on each side and these values were transformed to a log scale. Age and Genotype were analyzed using a two-way ANOVA with Tukey post-hoc tests.

***Novelty Suppressed Feeding (NSF)***

Testing was performed as previously described (2). Briefly, the NSF testing apparatus consisted of a plastic box (50 x 50 x 20 cm). The floor of which was covered with approximately 2 cm of wooden bedding and the arena was brightly lit (approximately 1000 lux). Mice were food restricted for 24 h prior to testing. At the time of testing, a single pellet of food (regular chow) was placed on a white paper platform positioned in the center of the box. Each animal was placed in a corner of the box, and a stopwatch was immediately started. The latency of the mice to begin eating in the arena was recorded. Immediately after the latency was recorded, the food pellet was removed from the arena. The mice were then placed back into their home cage. The latency to eat and the amount of food consumed in 5 min were measured (home cage consumption), followed by an assessment of post-restriction weight. A Kaplan-Meier survival analysis was used due to the lack of normal distribution of data. The Mantel-Cox log-rank test was used to evaluate differences between the experimental groups.

***Contextual Fear Conditioning (CFC)***

A 3-shock CFC paradigm was administered as previously described (4). CFC was conducted in chambers obtained from Actimetrics (Lafayette, IN), with internal dimensions of 7.4" L x 8.1" D x 7.9" H. The chambers had metal walls on each side, clear plastic front and back walls and ceilings, and stainless-steel bars on the floor. A house light (CM1820 bulb, 28v, 100mA) mounted directly above the chamber provided illumination. Each chamber was located inside a larger, insulated, plastic cabinet that provided protection from outside light and noise. Each cabinet contained a ventilation fan that was operated during the sessions. A paper towel dabbed with lemon solution was placed underneath the chamber floor. Mice were held outside the experimental room in their home cages prior to testing and transported to the conditioning apparatus individually in standard mouse cages. Chambers were cleaned with 70% EtOH after each run. Mice were placed into the conditioning chamber and received shocks at 180 s, 240 s, and 300 s (2 s duration, 0.75 mA). Fifteen seconds after the last shock, mice were removed from the chamber. Overall, the training session lasted 317 s. During re-exposure, mice were placed in the conditioning chamber for 3 minutes and did not receive any shocks. All sessions were scored for freezing using FreezeFrame4. Age and Genotype were analyzed using a repeated measures two-way ANOVA with Tukey post-hoc tests.

**Drugs**

***4-hydroxytamoxifen (4-OHT)***

Prior to CFC, mice were intraperitoneally (i.p.) injected with 2 mg of 4-OHT (Sigma, St Louis, MO). Four-OHT was dissolved into a 10% EtOH/90% corn oil solution by sonication (10 mg/ml)(5). After an injection, mice were dark housed for 3 days in order to limit other stimuli that might induce *Arc* expression(6).

***Methoxy-X04***

To confirm plaque pathology mice were injected i.p with 0.2 mL of Methoxy-X04 (Tocris Bioscience #4920, Minneapolis, MN) 24 hours prior to sacrifice. The 1 mg/mL solution was prepared by combining 45% propylene glycol, 45% 1X phosphate buffered saline (1X PBS), 10% dimethylsulfoxide (DMSO), and 10 mg of Methoxy-X04.

**Immunohistochemistry**

Mice were deeply anaesthetized with an i.p. injection of a (*R*,*S*)-ketamine/xylazine mixture (Ketaset III, Ketamine HCl injection, Fort Dodge Animal Health, Fort Dodge, Iowa 100 mg/kg / 10 mg/kg) 1 hour following CFC context re-exposure. Mice were transcardially perfused with 1X PBS followed by 4% paraformaldehyde (PFA). Brains were removed and stored in 4% PFA for 24 hours at 4°C followed by 72 hours in a 30% sucrose solution in 1X PBS before freezing in OCT. Using a cryostat at -22°C, the frozen brains were coronally sectioned at 100 µm. Slices were kept in 1X PBS with 0.2% sodium azide before staining. Slices were rinsed 3x in 1X PBS and dehydrated in 50% MeOH/1X PBS for 2 hours. Slices were then rinsed 3x in 1X PBS/0.2% Triton X (0.2% PBST) and blocked in 0.2% PBST/10% Dimethyl sulfoxide (DMSO)/6% normal donkey serum (NDS). Slices were washed 3x with 1X PBS/ 0.2% Tween-20 with 10 µg/ml heparin (PTwH) before incubation with the primary antibodies was performed at 4°C for 3 days (rabbit anti c-Fos, 1:5000, SySy, Göttingen, Germany, #156003; chicken anti-GFP, 1:500, Abcam, Cambridge, MA, #ab13970) in PTwH/ 5% DMSO/ 3% NDS. Sections were then washed 3x in PTwH before being placed in secondary antibodies at 4°C overnight (Alexa 647 conjugated donkey anti-rabbit, 1:500, Life Technologies, Eugene, OR, #1826679; Cy2 conjugated donkey anti-chicken IgG 1:500, Jackson ImmunoResearch, West Grove, PA, #703-

225-155) in PTwH/ 3% NDS. Sections were washed 3x in PTwH and then 3x in 1X PBS before being mounted on slides and coverslipped with DPX (Sigma Aldrich, Darmstadt, Germany).

**Confocal microscopy**

Fluorescent confocal micrographs were taken at 20x magnification with a Leica TCS SP8 MP microscope and with LAS X software. Images were scanned using overview software from Leica and captured at a z-increment of 3 µm.

**Cell Quantification**

***Automated cell counting***

Cells were automatically quantified in 3D using custom scripts in Fiji, as recently described (7). c-Fos$^+$ cells were identified by first passing the image through a bandpass filter in Fourier space, subtracting the background using a rolling ball algorithm, and identifying the cells using the 3D Local Maxima Fast Filter, 3D Spot Segmentation, and 3D Manager plugins in the 3D ImageJ suite (8). To maximize the precision of the automated counts, all segmented objects were filtered by size, shape, and intensity variation.

***Manual cell counting***

An investigator blind to treatment counted eYFP$^+$ and c-Fos$^+$ immunoreactive cells bilaterally in the granule cell layer (GCL) of the DG or in the pyramidal layer (PL) of CA3 and CA1 throughout the entire rostro-caudal axis of the hippocampus (HPC). Cells were counted bilaterally using Fiji and normalized to the area of the GCL or PL. The average eYFP$^+$ and c-Fos$^+$ cells per mm$^2$ are presented in **Figures 4** and **5**.

**Registration to an Anatomical Atlas**

Immunohistochemistry-labeled coronal brain sections were aligned to an anatomical atlas using the WholeBrain package in R (9). The atlas plate most closely corresponding to each section was identified using openbrainmap.org, and WholeBrain was used to automatically align the brain section to the corresponding atlas plate. All sections were manually curated to ensure an accurate fit, and when necessary, the alignment automatically generated by WholeBrain was manually adjusted. In some cases, due to uneven cutting or damage to the section, different hemispheres from the same section were aligned to different atlas plates. Misaligned or damaged regions were excluded from further analysis.

**Data Integration and Analysis**

Cell information, including location, intensity, and size, were imported into R from Fiji and copied into the WholeBrain object corresponding to the appropriate section. WholeBrain (9) was then used to convert the image coordinates of each cell into atlas coordinates and determine which brain region contained each cell. Data for each individual label were imported separately, but processed in parallel. Cells mapping to areas not expected to contain cells, such as fiber tracts and ventricles, or mapping outside the identified regions of the atlas, were excluded. Additionally, cells mapping to cortical layer 1 were also excluded. The area of each region was calculated using Gauss's area formula. Areas and cell counts were aggregated across layers to yield a single value for each region, and aggregate areas were converted to volumes and used to normalize the aggregate cell counts for each region. Normalized counts (cells per $mm^3$) were used in all further network analyses.

**Correlation and Network Analyses**

For c-$Fos^+$ network analysis, only common regions mapped across all experimental groups that were represented in a minimum n of 4 animals per group were included. Pearson correlations between regions were calculated using the Hmisc package in R, with pairwise removal of missing cases. Significance of pairwise regional correlation differences between experimental groups was calculated using permutation analysis. Group labels were randomly shuffled and correlations were recomputed 1000 times to generate a null distribution of the pairwise regional correlation differences between groups. Correlation differences were compared to these null distributions to determine the p-value. For the cluster network maps, relevant functional connections were retained by thresholding connections at $p < 0.05$. To ensure that both relevant positive and negative functional connections were used for community detection, the absolute Pearson values were used as edge weights. Using the igraph and tidygraph packages, the cluster fast greedy algorithm was used for community detection and visualized as a color-coded force-directed network (Fruchterman and Reingold layout) and as a dendrogram. Communities and nodes were color-coded and scales were changed for edge connections (correlation strength).

To compare global network properties of c-Fos$^+$ expression across experimental groups, networks were again constructed based on Pearson correlations and edges were thresholded at a p-value < 0.05. For each region, the clustering coefficient and measures of centrality, such as degree, betweenness centrality, and efficiency were calculated using the tidygraph and igraph packages. These measures were averaged across all regions to calculate global network statistics. To visualize only the strongest functional relationships, edges used for network plots (Fig 3A-D) were thresholded using a p-value of 0.01. Networks were visualized using ggraph and summary statistics were plotted in GraphPad Prism 9.

**Statistical Analysis**

All data were analyzed using JMP© software (SAS Institute, Cary, NC) and GraphPad Prism 9 (Boston, MA). Data were analyzed using two- or three-way ANOVA, with repeated measures when appropriate. Tukey or Bonferroni post-hoc tests were performed where appropriate. For the regression analysis, the average CFC freezing (%) versus marble burying (%) or co-labeled (%) for each individual mouse at 6 months of age was plotted. Alpha was set to 0.05 for all analyses. Data are expressed as means +/- SEM. All statistical tests and *p* values are included in the supplemental tables.

### HUMAN METHODS

#### Measuring NPIQ-Anxiety

To assess the degree of anxiety, the Neuropsychiatric Inventory brief Questionnaire (NPI-Q) form was used to assess neuropsychiatric symptoms (10). The NPI-Q is an informant-based, well-validated questionnaire, used widely in the research setting, which consists of the following 12 items: delusions, hallucinations, agitation/aggression, depression/dysphoria, euphoria/elation, anxiety, apathy, disinhibition, irritability/lability, aberrant motor behavior, sleep, and appetite/eating disorder. The NPI-Q includes one question for each of the 12 items, either yes or no (indicating the presence or absence of the symptom) and, if present, rated for severity as mild, moderate, and severe.

#### Cerebrospinal fluid and plasmatic protein levels

Lumbar puncture was performed with a 20- or 24-gauge spinal needle as described in the ADNI procedures manual (http://www.adni-info.org/). Briefly, CSF was collected into tubes provided to each site, then transferred into polypropylene transfer tubes followed by freezing on dry ice within 1 hour after collection, and shipped overnight to the ADNI Biomarker Core laboratory at the University of Pennsylvania Medical Center on dry ice. A more detailed description of the methodology regarding the sample acquisition, sample processing and analysis, as well as quality control procedures is available at the ADNI websites (http://adni.loni.usc.edu/methods/biomarker-analysis/proteomic-analysis and http://adni.loni.usc.edu/data-samples/biospecimen-data).

**ADNI dataset statistical analysis**

For descriptive statistics, mean and standard deviation and frequency and percent were reported for continuous and categorical variables, respectively. For group comparison, generalized linear model and chi-square test were used as appropriate. All statistical tests were adjusted for age, years of education, and sex. We considered the presence or absence of anxiety as the primary predictor and then evaluated anxiety severity as a secondary measure. Due to the low prevalence of the severe symptom group (n=11 out of n=1661), moderate and severe groups were combined.

First, the prevalence of anxiety was compared by baseline diagnosis group, and the interaction with sex was tested using ordinal logistic regression (Model 1). Second, among the mild cognitively impaired participants at baseline who were followed at least one time point and with CSF measures at baseline, Cox hazard regression examined the predictive utility of baseline anxiety for dementia transition in the amyloid positive participants (Model 2). To ensure consistency of the follow-up time across participants from different ADNI cohorts, 5-year dementia transition was examined. Sex interaction with anxiety and amyloid positivity were examined.

Finally, we compared the brain volume measures (68 bilateral cortical, 22 gray matter subcortical, and 5 corpus callosum ROIs) from Freesurfer by anxiety and tested the interactions between anxiety and sex using linear regression modeling. Among 1661 participants, 11 subjects were excluded due to bad image quality, resulting in 1650 subjects. All models were adjusted for estimated intracranial volume, age, sex, and education. We performed multiple comparison correction controlling for false discovery rate (11).

### REFERENCES


1. Mastrodonato A, Martinez R, Pavlova IP, LaGamma CT, Brachman RA, Robison AJ, Denny CA (2018): Ventral CA3 Activation Mediates Prophylactic Ketamine Efficacy Against Stress-Induced Depressive-like Behavior. *Biol Psychiatry* 84: 846–856.
2. Brachman RA, McGowan JC, Perusini JN, Lim SC, Pham TH, Faye C, *et al.* (2016): Ketamine as a Prophylactic Against Stress-Induced Depressive-like Behavior. *Biol Psychiatry* 79: 776–786.
3. Walf AA, Frye CA (2007): The use of the elevated plus maze as an assay of anxiety-related behavior in rodents. *Nat Protoc* 2: 322.
4. Perusini JN, Cajigas SA, Cohensedgh O, Lim SC, Pavlova IP, Donaldson ZR, Denny CA (2017): Optogenetic stimulation of dentate gyrus engrams restores memory in Alzheimer's disease mice. *Hippocampus* 27: 1110–1122.
5. Cazzulino AS, Martinez R, Tomm NK, Denny CA (2016): Improved specificity of hippocampal memory trace labeling. *Hippocampus* 26: 752.
6. Denny CA, Kheirbek MA, Alba EL, Tanaka KF, Brachman RA, Laughman KB, *et al.* (2014): Hippocampal memory traces are differentially modulated by experience, time, and adult neurogenesis. *Neuron* 83: 189–201.
7. Leal Santos S, Stackmann M, Zamora AM, Mastrodonato A, De Landri A V., Vaughan N, *et al.* (2021): Propranolol decreases fear expression by modulating fear memory traces. *Biol Psychiatry*. https://doi.org/10.1016/j.biopsych.2021.01.005
8. Schindelin J, Arganda-Carreras I, Frise E, Kaynig V, Longair M, Pietzsch T, *et al.* (2012, July): Fiji: An open-source platform for biological-image analysis. *Nature Methods*, vol. 9. Nat Methods, pp 676–682.
9. Fürth D, Vaissière T, Tzortzi O, Xuan Y, Märtin A, Lazaridis I, *et al.* (2018): An interactive framework for whole-brain maps at cellular resolution. *Nat Neurosci* 21: 139–153.
10. Kaufer DI, Cummings JL, Ketchel P, Smith V, MacMillan A, Shelley T, *et al.* (2000): Validation of the NPI-Q, a brief clinical form of the Neuropsychiatric Inventory. *J Neuropsychiatry Clin Neurosci* 12: 233–239.
11. Benjamini Y, Hochberg Y (1995): Controlling the False Discovery Rate: A Practical and Powerful Approach to Multiple Testing. *Journal of the Royal Statistical Society: Series B (Methodological)* 57: 289–300.

### SUPPLEMENTARY FIGURES AND FIGURE LEGENDS

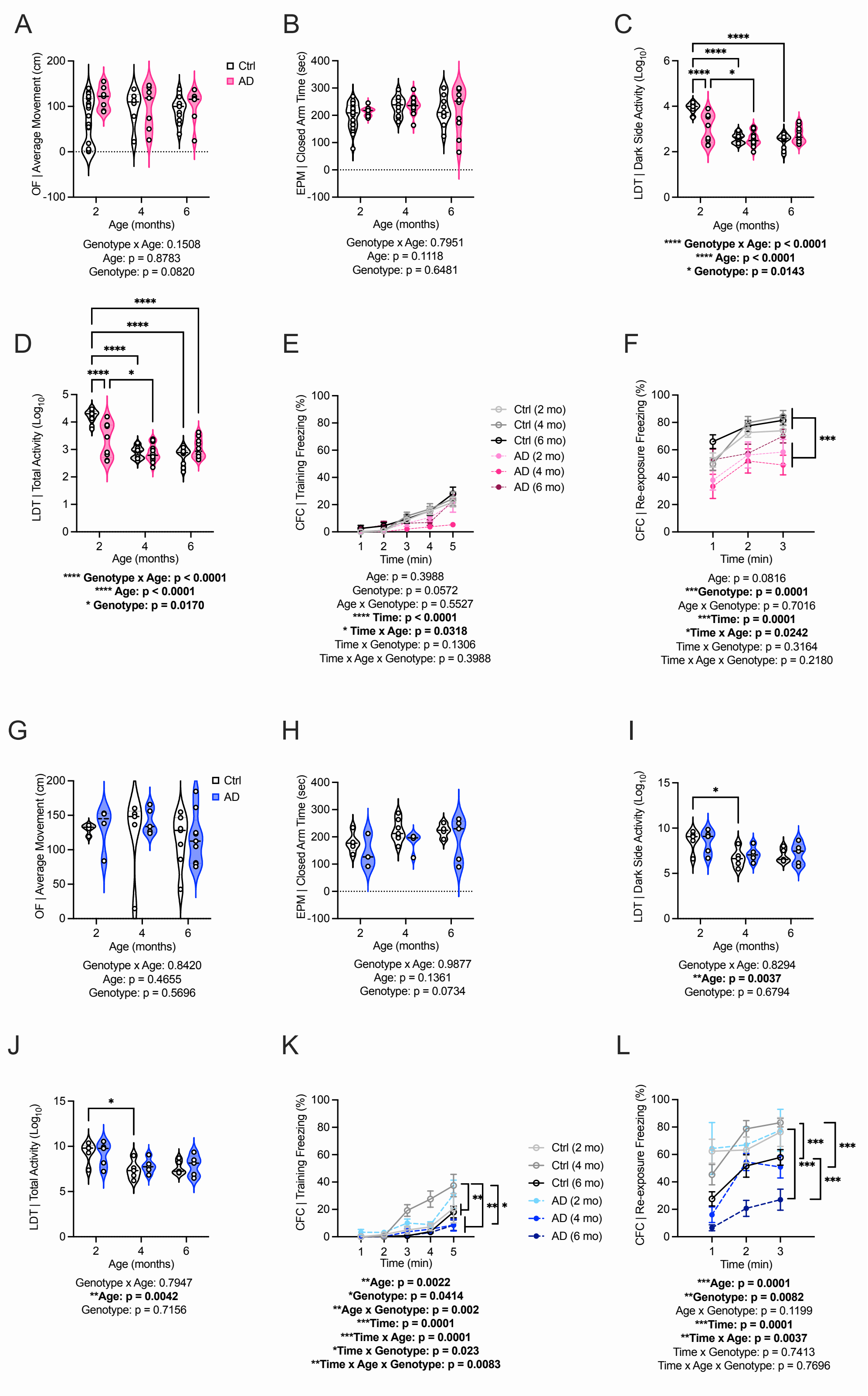

**Supplemental Fig. 1.** *Avoidance behavior and cognitive impairment in aging and AD mice.* (**A**) Female mice exhibit similar average movement in the OF across ages. **(B)** Closed arm time was similar among female groups. (**C-D**) In the LDT, female AD mice exhibit a decrease in dark side and total activity starting at 2 months of age compared to female Ctrl mice. This activity declines with age. (**E-F**) Line graphs representing freezing behavior during training and re-exposure. During CFC training, there is a Time x Age significant interaction. (**F**) However, during context re-exposure, female AD mice exhibit decreased freezing starting at 2 months of age compared to female Ctrl mice. (**G**) Male mice exhibit comparable average movements in the OF across ages. (**H**) In the EPM, time spent in the closed arms is comparable in male groups. (**I-J**) In the LDT, there is a significant effect of Age on dark side and overall activity in the male groups. **(K)** During CFC training, there is a Time x Age x Genotype significant interaction during in male mice where 4-month male Ctrl mice exhibit increased freezing with shock. (**L**) During context re-exposure, at 6 months of age, male AD mice exhibit decreased freezing compared to male Ctrl mice. (n=4-18 mice per group). Error bars represent ± SEM. *$p<0.05$, **$p<0.01$, ***$p<0.001$. OF, open field; EPM, elevated plus maze; LDT, light dark test; CFC, contextual fear conditioning; mo, months; cm, centimeters; Ctrl, control; AD, Alzheimer's disease.

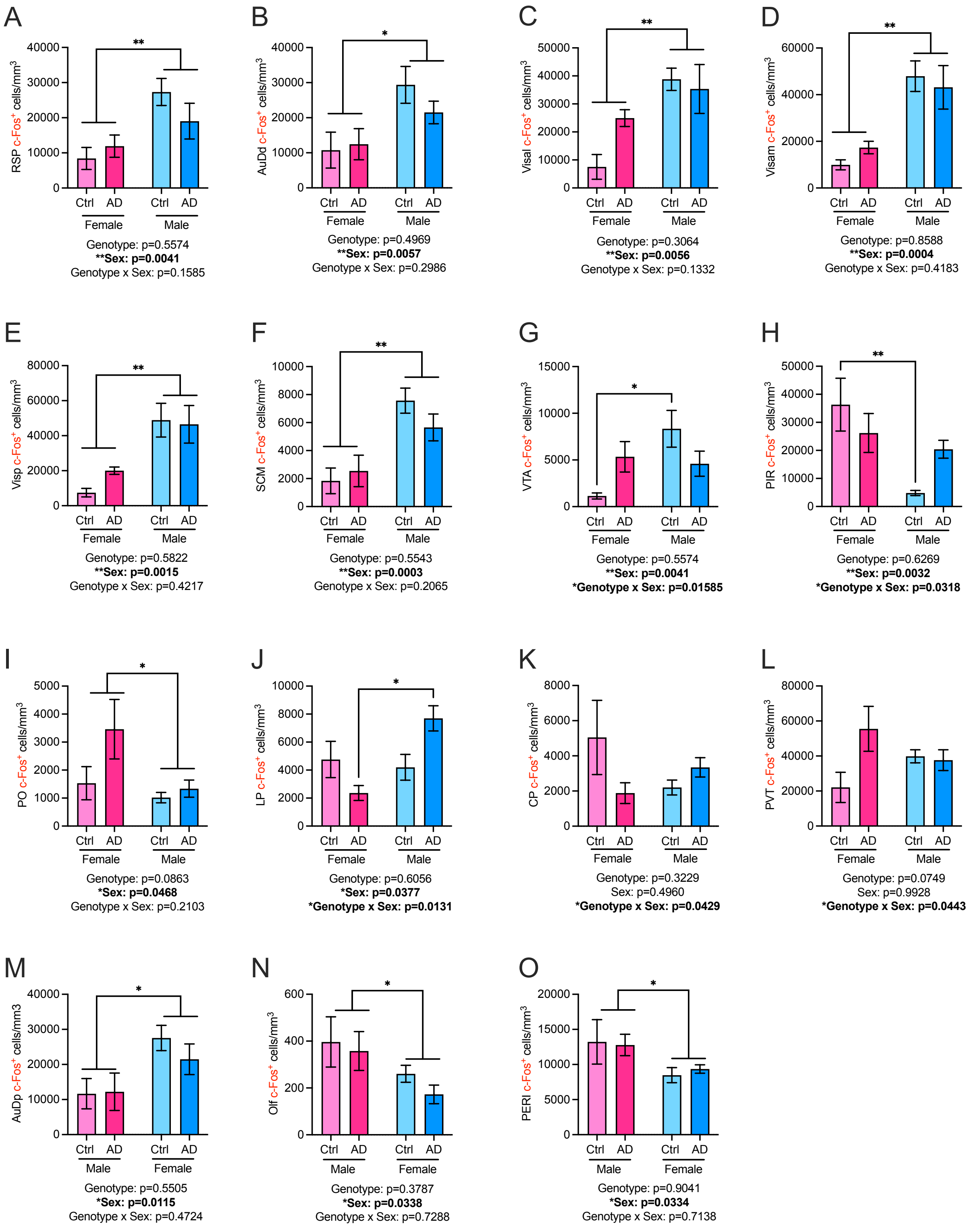


**Supplemental Fig. 2.** *Brain-wide c-Fos$^{+}$ activation in male and female mice.*

(**A-F**) There was a main effect of Sex in the RSP, AuDd, Visal, Visam, Visp, and SCM; male mice exhibited greater c-Fos+ expression when compared with female mice. (**G**) In the VTA, there was a significant effect of Sex and a Genotype x Sex interaction. Male Ctrl mice exhibited increased c-Fos+ expression when compared with female Ctrl mice. (**H**) In the PIR, there was a significant effect of Sex and a Genotype x Sex interaction. Male Ctrl mice exhibited less c-Fos+ expression when compared with female Ctrl mice. (**I**) In the PO, there was a main effect of Sex; male mice exhibited less c-Fos+ expression when compared with female mice. **(J**) In the LP, there was a significant effect of Sex and a Genotype x Sex interaction. Male Ctrl mice exhibited increased c-Fos+ expression when compared with female Ctrl mice. (**K-L**) In the CP and PVT, there was a significant Genotype x Sex interaction, but no post-hoc tests reached significance. (**M-O**) In the AuDp, Off, and PERI, there was a main effect of Sex. In the AuDp, male mice exhibited increased c-Fos+ expression when compared with female Ctrl mice. In the Off and PERI, male mice exhibited decreased c-Fos+ expression when compared with female Ctrl mice. (n=4-7 mice per group). Error bars represent ± SEM. *p<0.05, **p<0.01, ***p<0.001. Ctrl, control; AD, Alzheimer's disease; RSP, retrosplenial area; AudD, dorsal auditory area; Visal, anterolateral visual area; Visam, anteromedial visual area; Visp, primary visual area; SCM, superior colliculus; VTA, ventral tegmental area; PIR, piriform area; PO, posterior complex of the thalamus; LP, lateral posterior nucelus of the thalmus; CP, caudoputamen; PVT, paraventricular nucleus of the thalamus; Audp, primary auditory area; OLF, olfactory areas; PERI, perirhinal area.

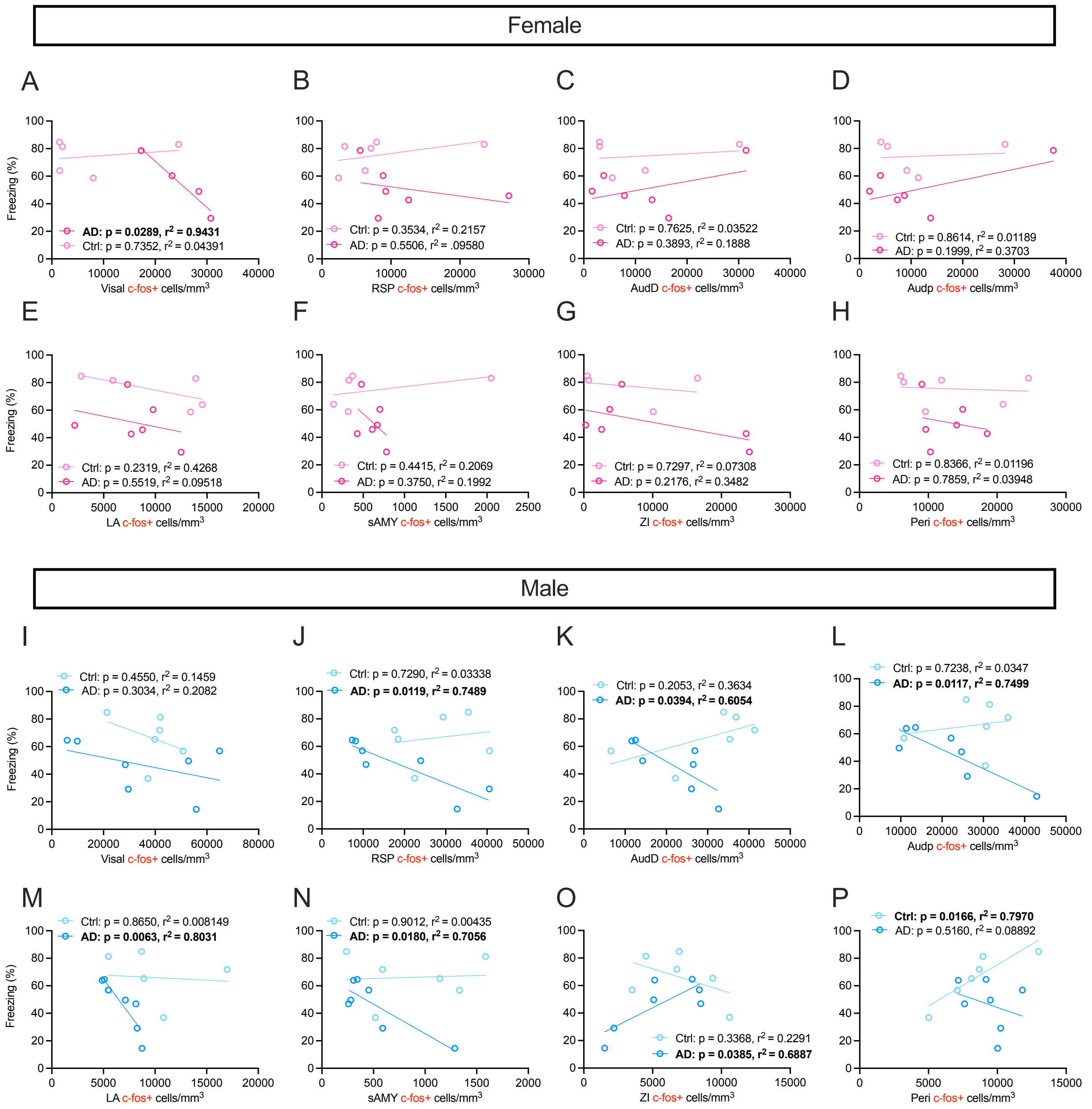


**Supplemental Fig. 3.** *Correlations between c-Fos and freezing behavior.*

(**A**) The number of Visal c-Fos$^+$ cells were negatively correlated with freezing (%) in AD female mice only. (**B-H**) No significant correlations between c-Fos$^+$ expression in identified brain regions and freezing (%) were observed in female mice. (**I**) No significant correlations were observed between the number of Visal c-Fos$^+$ cells and freezing (%) in male mice. (**J-N**) c-Fos$^+$ expression in the RSP, AudD, Audp, LA, and sAMY regions was negatively correlated with freezing (%) in male AD mice. (**O**) c-Fos$^+$ expression in the ZI was positively correlated with freezing (%) in male AD mice. (**P**) c-Fos$^+$ expression in the Peri was positively correlated with freezing (%) in male Ctrl mice. (n=4-7 mice per group). *p<0.05, **p<0.01, ***p<0.001. Ctrl, control; AD, Alzheimer's disease; Visal, anterolateral visual area; RSP, retrosplenial area; AudD, dorsal auditory area; Audp, primary auditory area; LA, lateral amygdala; sAMY, striatum like amygdalar nuclei; ZI, zona incerta; PERI, perirhinal area.

**SUPPLEMENTARY TABLES**

**Table S1.** *Statistics for mouse behavioral data*. This table lists all statistics for female and male behavioral data in **Figure 1** and Supplemental **Figures 1** and **2**.

**Table S2.** *Network analysis*. This table includes the statistics for the heatmaps and network analysis in **Figure 2** and Supplemental **Figures 2** and **3**.

**Table S3.** *Engram cell count statistics*. This table lists the statistics used for the encoding, retrieval, and co-labeled cell count data in **Figure 3**.

**Table S4.** *ADNI demographics.* This table outlines the demographics of the Alzheimer's disease neuroimaging initiative (ADNI) cohort.

**Table S5.** *ADNI anxiety and amyloid measures*. This table lists the statistics for anxiety measures in the ADNI cohort in **Figure 6**.

**Table S6.** *ADNI odds ratio*. This table lists statistics for the odds of transitioning to dementia based on anxiety measures represented in the **Figure 6**.

**Table S7.** *ADNI estimated survival probability*. This table includes all of the raw data from the survival analysis in **Figure 6** with 95% CIs for each stratum (Anxiety, Abeta positivity and gender).

**Table S8.** *ADNI survival analysis bar graph*. This table includes the statistics from the survival analysis in bar graph form in **Figure 6**.

**Table S9.** *ADNI brain volumes and anxiety*. This tables provides the prediction analysis model and results shown in **Figure 6**.

**Table S10.** *ADNI Predictors of dementia transition***.** This table includes statistics for anxiety and its relation to brain volume in both male and female subjects in **Figure 7**.
